# Structural basis of iron piracy by human gut *Bacteroides*

**DOI:** 10.1101/2024.04.15.589501

**Authors:** Augustinas Silale, Yung Li Soo, Hannah Mark, Rachel N. Motz, Arnaud Baslé, Elizabeth M. Nolan, Bert van den Berg

**Affiliations:** Biosciences Institute, Newcastle University, Framlington Place, Newcastle upon Tyne, NE2 4HH, United Kingdom; Department of Chemistry, Massachusetts Institute of Technology, Cambridge, Massachusetts 02139, United States

## Abstract

Iron is an essential element that can be growth-limiting in microbial communities, particularly those present within host organisms. To acquire iron, many bacteria secrete siderophores, secondary metabolites that chelate ferric iron. These iron chelates can be transported back into the cell via TonB-dependent transporters in the outer membrane, followed by intracellular liberation of the iron. Pathogenic *Escherichia coli* and *Salmonella* produce siderophores during gut infection. In response to iron starvation, the human gut symbiont *Bacteroides thetaiotaomicron* upregulates an iron piracy system, XusABC, which steals iron-bound siderophores from the invading pathogens. Here, we investigated the molecular details of xenosiderophore uptake across the outer membrane by the XusAB complex. Our crystal and cryogenic electron microscopy structures explain how the XusB lipoprotein recognises iron-bound xenosiderophores and passes them on to the XusA TonB-dependent transporter. Moreover, we show that Xus homologues can transport a variety of siderophores with different iron-chelating functional groups.

## Introduction

Iron is essential for most organisms, including almost all bacteria. It is a protein cofactor required for enzymatic reactions and electron transport during various cellular processes. While iron is one of the most abundant transition metals on Earth, its bioavailability is limited due to the extremely low solubility (∼10^−18^ M) of its predominant ferric (Fe^3+^) form at physiological pH (*1*). Many Gram-negative bacteria release siderophores, iron-chelating secondary metabolites, into the environment which, once bound to ferric iron, can be transported back into the cell via TonB-dependent transporters in the outer membrane (OM) (*2*). A classic example is enterobactin (Ent), which is produced by Enterobacteriaceae and consists of three iron-chelating catecholate groups connected via amide linkers to a cyclic triserine lactone (*3*, *4*). Ferric enterobactin (FeEnt) is taken up by the FepA and IroN transporters in *E. coli* and *Salmonella* (*5–7*), and iron is liberated from the FeEnt complex inside the cell (*8*, *9*). Siderophore-mediated iron scavenging is important for pathogens during infection, when the host restricts iron availability to starve invading bacteria via nutritional immunity (*10–12*). Additionally, secreted siderophores are not necessarily taken up by the same bacterium that produced them, resulting in complex iron availability-dependent interactions in microbial communities (*13*).

The gastrointestinal tract is home to a diverse community of microorganisms that cooperate and compete for available nutrients both with the host and between themselves (*14*, *15*). Depriving the gut microbiota of iron eventually results in irreversible structural changes in gut microbial communities (*16*, *17*). Bacteroidota, the dominant phylum of diderm bacteria in the distal gut, are not known to produce siderophores. *Bacteroides thetaiotaomicron* (*B. theta*) has been shown to prefer heme as an iron source over soluble ferrous iron (*18*). *B. theta* likely acquires heme from the diet (*19*) as well as from dead intestinal cells and dead bacteria. Iron-bound siderophores produced by commensal bacteria and fungi as well as various pathogens are another source of bioavailable iron for gut bacteria that have transporters to take them up (*20*). It was recently shown that during *Salmonella* infection in the mouse gut, when iron availability becomes limiting, *B. theta* upregulates a novel iron uptake system that steals iron-bound siderophores produced by the pathogenic *Salmonella* (*21*). This <u>x</u>enosiderophore (i.e., foreign siderophore) <u>u</u>tilization <u>s</u>ystem (Xus) consists of the TonB-dependent transporter XusA, the surface-exposed lipoprotein XusB and the PepSY domain-containing inner membrane protein XusC (*21*, *22*). The XusABC system provides resilience to *B. theta* during infection, but XusB secreted by *B. theta* and bound to xenosiderophores also acts as an iron reservoir for the pathogen (*21*, *22*). It is therefore important to understand the molecular details of this iron piracy system to obtain a deeper understanding of pathogen-symbiont interactions inside the gut.

Here, we present crystal structures of the apo- and xenosiderophore-bound XusB lipoproteins from several Bacteroidota species, which reveal the mechanism of xenosiderophore capture. We also use single particle cryo-EM to investigate how XusB interacts with the XusA TonB-dependent transporter. In vitro binding studies and structural bioinformatics analyses reveal distinct subclasses of xenosiderophore utilization systems in Bacteroidota. Together, our results provide mechanistic insights into xenosiderophore uptake across the OM of the gut commensal *B. theta* and the pathobiont *Bacteroides fragilis*.

## Results

### Crystal structures of apo and xenosiderophore-bound *B. theta* XusB

We expressed recombinant XusB of *B. theta* (BtXusB; UniProt accession Q8A622) lacking the signal sequence and the lipid anchor cysteine in *E. coli* BL21(DE3) and determined its crystal structure using data to 1.56 Å resolution (Fig. 1A and B, Table S1). BtXusB has a seven-bladed β-propeller fold with a distinct 15-residue loop, which we termed the hook, inserted into the β2 blade and protruding outwards from the β-propeller (Fig. 1A). The β-propeller pocket (Fig. 1B) has been suggested to bind FeEnt based on recent computational docking (*22*).

**Fig. 1.**
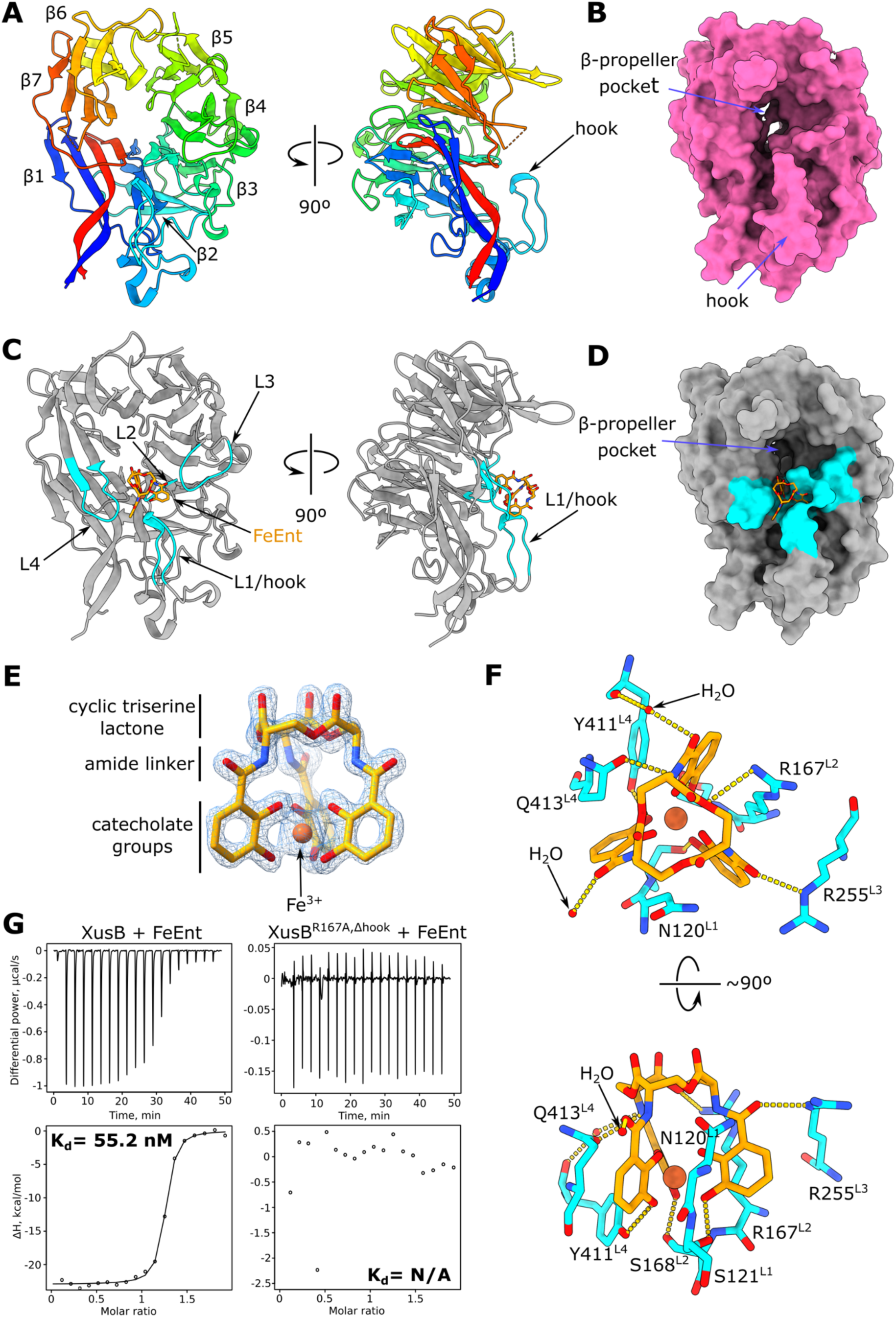
BtXusB binding to FeEnt. (**A**) Crystal structure of apo BtXusB at 1.56 Å. The cartoon is coloured in rainbow; the N-terminus is blue, the C-terminus is red. Blades of the β-propeller (β1-7) are labelled starting from the N-terminus. (**B**) Surface representation of apo BtXusB. (**C**) Co-crystal structure of BtXusB bound to FeEnt at 1.5 Å. FeEnt is depicted as an orange stick model. The four FeEnt binding loops (L1-L4) are shown in cyan. (**D**) Surface representation of BtXusB bound to FeEnt. (**E**) FeEnt model fit into the 2mF_o_-DF_c_ electron density map at 2σ. (**F**) BtXusB residues interacting with FeEnt. The yellow dashed lines show likely hydrogen bonds. (**G**) Representative ITC experiments where 250 μM FeEnt was titrated into 25 μM BtXusB (n=4 experiments) or 25 μM BtXusB^R167A,Δhook^ variant (n=2 experiments). Integrated heats were fitted to a single binding site model, giving the apparent K_d_ value for the XusB-FeEnt titration. No K_d_ value could be determined for the XusB^R167A,Δhook^ titration.

We co-crystallized recombinant BtXusB with FeEnt and determined the crystal structure of the complex to 1.50 Å. BtXusB binds FeEnt via four loops (Fig. 1C), including the hook. Interestingly, FeEnt binds off the β-propeller pocket rather than in it, and the distance to the previously modelled binding site is ∼10 Å (Fig. 1D) (*22*), underscoring the importance and value of experimental protein-ligand structure determination. The FeEnt electron density in the co-crystal structure is of very high quality (Fig. 1E), with the cyclic triserine lactone, amide linkers and Fe^3+^-ligating catecholate groups fully resolved. The catecholate arms of FeEnt are not perpendicular to the triserine lactone ring plane but slanted to one side (Fig. 1E, Movie S1), in agreement with the structure of FeEnt in isolation (*4*). FeEnt does not undergo conformational changes upon binding to BtXusB. Instead, BtXusB loops close in on FeEnt, forming a 419 Å^2^ interaction interface (Fig. S1 and S2, Movie S2). All three catecholate arms of FeEnt interact with BtXusB loops L1-4 (Fig. 1C and F). Sidechains of N120, R167 and Q413 slot between the slanted catecholate arms, while S121, S168 and Y411 sidechains form hydrogen bonds with the *meta* oxygens of the catecholate groups (Fig. 1F, Movie S1). One amide linker of FeEnt hydrogen-bonds to a single water molecule, another to a water molecule and the sidechain of Q413, and the third to the sidechain of R255 (Fig. 1F). The sidechain of R167 forms a hydrogen bond with the triserine lactone ring—the only interaction between BtXusB and this part of FeEnt.

BtXusB binds FeEnt with a dissociation constant value of ∼55 nM as determined by isothermal titration calorimetry (ITC) (Fig. 1G). We constructed a BtXusB variant, BtXusB^R167A,Δhook^, with the R167A substitution and deletion of residues 119-122 which form the tip of the hook. Titration of FeEnt into BtXusB^R167A,Δhook^ resulted in reduced injection heats and no saturation (Fig. 1G), which we interpret as lack of binding. The ITC results strongly suggest that the binding site observed in the co-crystal structure is the only FeEnt binding site on BtXusB.

Ent secreted by pathogens and commensals can be taken up by any other Gram-negative bacterium that expresses suitable TonB-dependent transporters, such as FepA and XusA (*5*, *21*). Furthermore, as part of the innate immune response host cells secrete lipocalin-2, which sequesters Ent and deprives pathogens of iron (*23*, *24*). Therefore, production of Ent by the pathogen can be less effective during infection. *Salmonella* and some *E. coli*, e.g. many uropathogenic strains (*25*), have in turn evolved a strategy to prevent waste of resources and secure iron by making chemically modified versions of Ent that require the TonB-dependent transporter IroN for import and do not bind to lipocalin-2, such as di-*C*-glucosylenterobactin (DGE), also known as salmochelin S4 (*26*, *27*). DGE has the same core structure as Ent, with two of the catechol groups *C-*glucosylated at the C5 position (Fig. 2A). Notably, *E. coli* FepA does not import DGE (*7*). We soaked apo BtXusB crystals with iron-bound DGE (FeDGE) and determined the crystal structure of FeDGE-bound BtXusB to 1.80 Å resolution (Fig. 2B). One of two protein chains in the asymmetric unit had FeDGE bound, which we could confidently build into the ligand density (Fig. 2B,C). BtXusB binds FeDGE via the same loops as FeEnt with additional interactions with the two glucosyl modifications, Glc1 and Glc2 (Fig. 2D). Glc1 hydroxyl groups hydrogen-bond to two water molecules and the hook loop via the backbone carbonyl oxygen of N120 and the side chain of S123. Glc2 contacts the hook via the sidechain of S122, L2 via the backbone of R167, and L3 via the backbone of S254. Additionally, Glc2 interacts with a single water molecule and the hydroxyl group of Y189, which is not part of the four loops that bind FeEnt. The electron density for Glc1 was weaker than for Glc2 (Fig. 2C), which suggests that Glc1 is bound less tightly than Glc2. The two glucosyl modifications slot between the ligand-binding loops (Fig. 2E). One notable difference between the FeEnt- and FeDGE-bound structures is that the amide linker of one of the catechol arms of FeDGE does not hydrogen-bond with the sidechain of L3 R255 as observed for FeEnt (Fig. 2F). This rearrangement is likely the result of the C6 atom of Glc2 nudging the sidechain of S254 by ∼1.7 Å and pushing the entire L3 further away from the siderophore. Consequently, the sidechain of R255 swings away by ∼5 Å and can no longer contact the amide linker. ITC data suggest that the amide-R255 interaction is important for tight xenosiderophore binding, as the FeDGE-BtXusB interaction has a ∼2-3-fold lower apparent dissociation constant value compared to FeEnt (Fig. 2G and Table S2), even though the glucosyl modifications of FeDGE form additional contacts with BtXusB. In addition, the decreased mobility of the constrained Glc modifications likely results in an entropic penalty for bound FeDGE. ITC data fitting results support this prediction, as the BtXusB-FeDGE interaction has an estimated -TΔS value ∼2.3 kcal/mol smaller than the BtXusB-FeEnt interaction, while the ΔG terms are similar for both with a difference of ∼0.6 kcal/mol (Table S2).

**Fig. 2.**
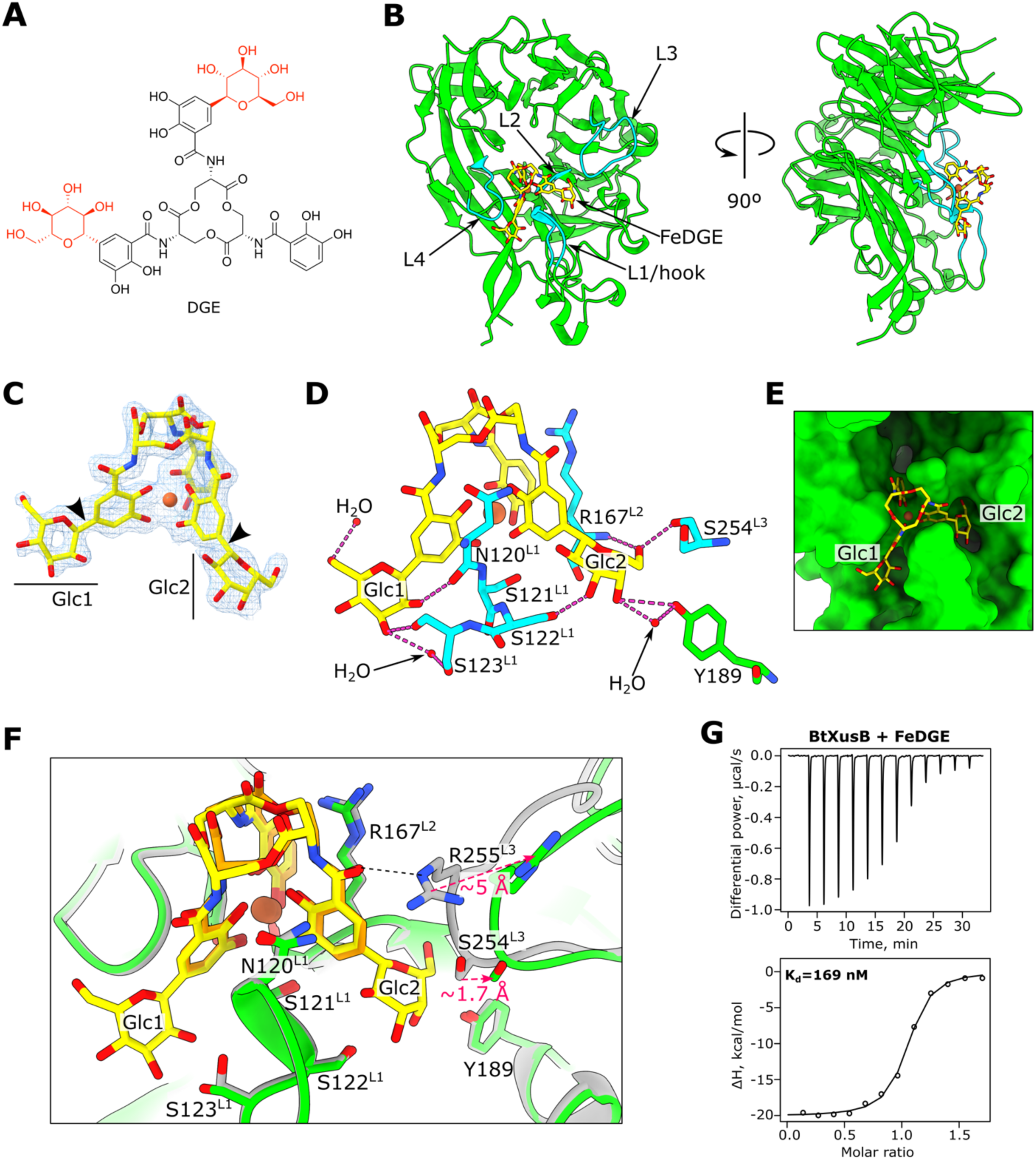
BtXusB binding to FeDGE. (**A**) Chemical structure of DGE. The glucosyl groups are in red. (**B**) Crystal structure of BtXusB (green) bound to FeDGE (yellow). The siderophore-binding loops are shown in cyan. (**C**) FeDGE model fit into the 2mF_o_-DF_c_ electron density map at 1σ. The arrowheads point to the C-C bonds between the catechol groups and the C1 atoms of the glucosyl moieties. (**D**) Hydrogen bonding network formed between the glucosyl groups of FeDGE and BtXusB. (**E**) The Glc1 and Glc2 groups of FeDGE occupy slots between the xenosiderophore-binding loops of BtXusB (green surface). (**F)** Superposition of BtXusB-FeEnt (grey and orange, respectively) and BtXusB-FeDGE (green and yellow, respectively) crystal structures. Hook loop residues N120-S123 and L2 R167 are in almost identical conformations in the two structures, but L3 is pushed away in the FeDGE structure due a clash between Glc2 of FeDGE and S254 of L3. The black dashed line indicates the hydrogen bond between R255 and the amide carbonyl of FeEnt. (**G**) Representative ITC experiment where 154 μM FeDGE was titrated into 17.3 μM BtXusB (n=2 experiments). Integrated heats were fitted to a single binding site model, giving the apparent K_d_ value.

### Cryo-EM structure of native XusAB complex from *B. theta*

FeEnt is sufficient to rescue growth of *B. theta* under iron limiting conditions (Fig. S3). We reasoned that the proteins encoded by the Xus operon would be expressed to high enough copy number for purification and structural characterisation if the cells were cultured under iron-limiting conditions. We purified the native XusAB complex from the *B. theta bt2064-his* strain grown in minimal medium supplemented with the iron chelator bathophenanthroline disulfonate (BPS) (Fig. S3, Methods). We determined the structure of XusAB by single particle cryo-EM to a global resolution of 2.7 Å, with local resolution extending to 2.2 Å (Fig. 3A, Fig. S4, Table S3). The resolved portion of XusA has a classic TonB-dependent transporter fold: a micelle-embedded 22-strand β-barrel occluded by an N-terminal plug domain (*2*). The XusA extreme N-terminal carboxypeptidase-like domain of unknown function was not resolved in our structure likely because it is connected to the barrel via a flexible linker. The conformation of XusB in the XusAB complex is almost identical to that observed in the apo XusB crystal structure, with Cα-Cα RMSD = 0.9 Å. We did not observe any alternative conformations, sub- complexes or movement of XusA and XusB in the cryo-EM data.

**Fig. 3.**
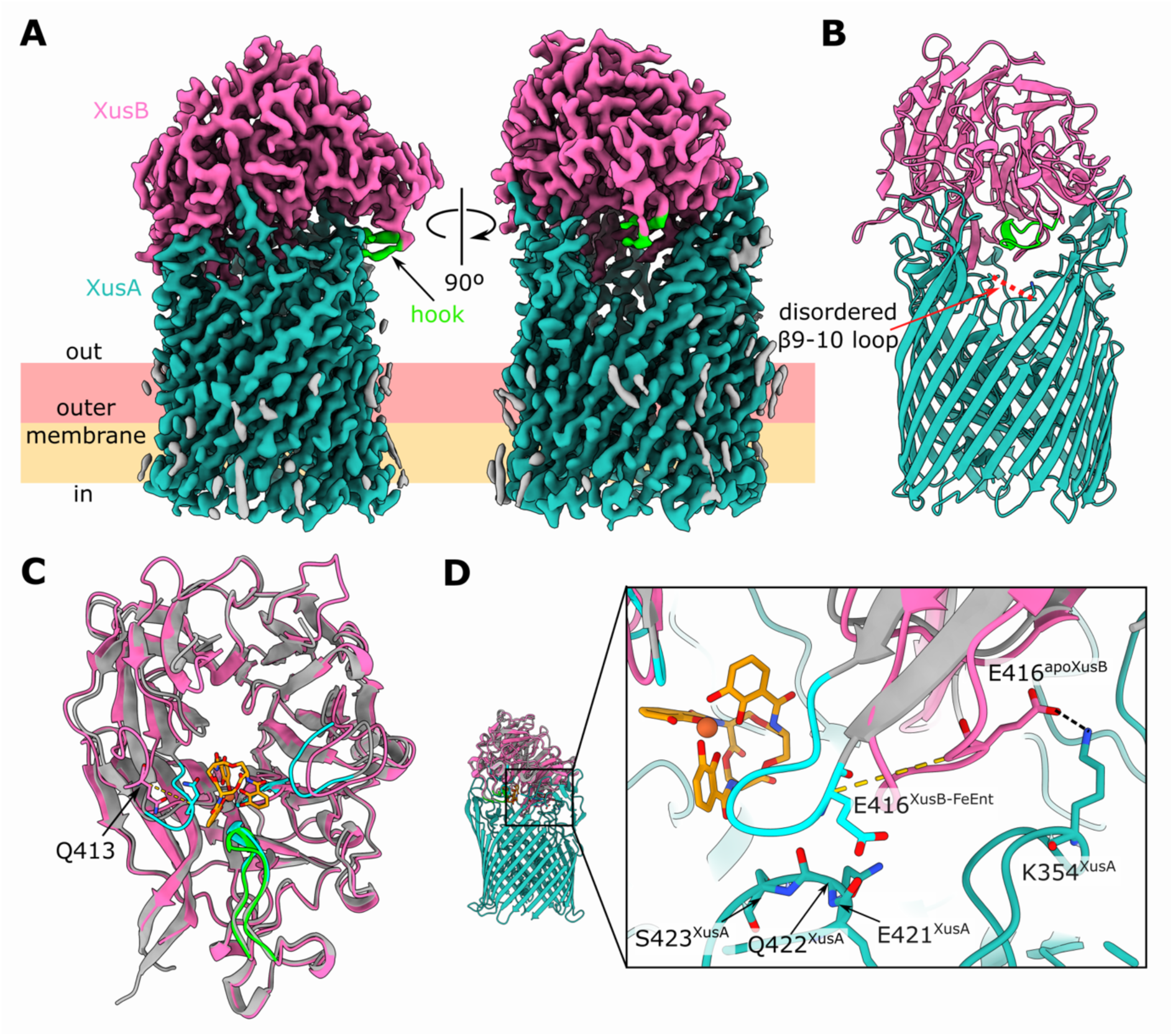
Cryo-EM structure of the native XusAB complex from *B. theta*. (**A**) Single particle cryo-EM reconstruction of the XusAB complex at 2.7 Å global resolution. (**B**) Model built into the cryo-EM density in cartoon representation. The red dashed line depicts the disordered extracellular loop between β strands 9 and 10 of the XusA β-barrel, corresponding to residues 464-473. (**C**) Structural alignment of the apo-XusB structure observed in the cryo-EM reconstruction (hot pink, hook in green) and the XusB-FeEnt co-crystal structure (grey, FeEnt binding loops in cyan). Cα-Cα RMSD = 1.6 Å. The dashed yellow line corresponds to a distance of 7.7 Å between the Cα atoms of Q413, which is part of the L4 FeEnt binding loop, observed in the two structures. (**D**) Close-up view of XusB L4 in the apo cryo-EM structure and the XusB-FeEnt co-crystal structure, viewed from a different orientation compared to (**C**). Colouring as in (**C**), XusA is in light sea green. The yellow dashes correspond to a distance of 11.3 Å between the Cα atoms of E416 in the two structures. The black dashes show a salt bridge between K354 of XusA and E416 of XusB in the cryo-EM structure. Residues E421, Q422 and S423 are part of the β7-8 extracellular loop of XusA.

XusB sits on top of the extracellular side of XusA like a lid, reminiscent of *B. theta* TonB-dependent transporter and surface-exposed lipoprotein complexes that take up glycans and vitamin B_12_ (*28*, *29*). The interaction interface of XusA and XusB is extensive: PISA analysis indicates 52 hydrogen bonds and 6 salt bridges, with a total interaction surface area of 2,795 Å^2^ (Fig. S5). XusB interacts with every extracellular loop of XusA, except for the loop between barrel strands β9 and β10, which is disordered (Fig. 3B). Although we did not observe FeEnt in the cryo-EM structure, the XusB hook facing towards a cavity enclosed by the XusAB complex (Fig. 3A,B) indicates that FeEnt binds inside this cavity. We speculate that XusB moves in a hinge-like motion opening and closing the XusAB cavity like a lid, thus transiently allowing extracellular xenosiderophores access to their binding site on the inward-facing side of XusB.

XusB binds FeEnt with high affinity (Fig. 1G), but FeEnt must somehow be transferred from XusB to XusA to achieve transport across the OM. Structural alignment of the cryo-EM XusB structure and the XusB-FeEnt co-crystal structure reveals that Q413, part of L4, must undergo a shift of 7.7 Å to interact with FeEnt (Fig. 3C). However, the position of L4 in the XusB-FeEnt structure would result in clashes with the XusA β7-8 extracellular loop within the apo complex (Fig. 3D). Furthermore, L4 in the apo complex is stabilised via a salt bridge between E416 and K354 of XusA β5-6 extracellular loop. This suggests that the FeEnt-bound state of XusB when in complex with XusA might be short-lived despite the high affinity of XusB for FeEnt. We speculate that the XusA β7-8 extracellular loop displaces L4 of XusB, aided by formation of the E416-K354 salt bridge, thus disrupting the FeEnt binding pocket and releasing the xenosiderophore to diffuse towards XusA, which would transport it across the OM. We envisage that the transfer of FeDGE from XusB to XusA proceeds via an identical mechanism to FeEnt.

We observed unexplained density extending from the sidechain of T401, which is part of the XusA β7-8 extracellular loop (Fig. S6). Together with flanking residues this threonine forms a DTA sequence, which matches the Bacteroidota O-glycosylation motif (*30*). We therefore conclude that the cryo-EM density extending from T401 corresponds to a glycan chain. Four sugar units can be discerned, including the branching deoxyhexose previously identified in the *B. fragilis* O-glycan (*31*). The T401 glycosylation site is located near residues 421-423 which are implicated in FeEnt release from XusB (Fig. 3D and Fig. S6). The sidechain of T401 and the O-glycan face the solvent rather than XusB, but we cannot rule out that the glycan modification affects the conformation of neighbouring XusA extracellular loops and that it could be functionally important.

### Structural variety of XusB homologues

BtXusB is annotated as a DUF4374-containing protein in UniProt (*32*), which has a β-propeller fold as demonstrated by our structural data. We investigated the predicted structures of DUF4374-containing homologues. The top hits from protein BLAST searches of the BtXusB sequence against Bacteroidota genomes have a sequence identity of around 60%. The reason for the relatively low sequence identity for the top BLAST hits is that the BtXusB hook position inside the β2 propeller blade is unique (Fig. 4A). Other Bacteroidota XusB homologues, such as *Parabacteroides distasonis* BDI_3402 (59.5% identity; UniProt accession A6LHE0) and *Barnesiella viscericola* BARVI_05925 (43.4% identity; UniProt accession W0ET74), are predicted to have their hook inserted in the β4 blade or, in the case of *B. fragilis* BF9343_4228 (20.2% identity; UniProt accession Q5L7E3), do not appear to have a hook at all (Fig. 4B-D and Fig. S7). These DUF4374-containing proteins are likely part of genuine xenosiderophore utilization systems, rather than performing some other function. They all contain a lipoprotein export signal that directs proteins to the outer leaflet of the OM (*33*). Additionally, they are encoded next to TonB-dependent transporters and PepSY domain-containing inner membrane proteins that likely reduce siderophore-bound ferric iron to facilitate dissociation of the iron-siderophore complex (*34*) (Fig. 4E). *P. distasonis* also encodes a periplasmic esterase (BDI_3403) in the same operon as the XusB homologue which might liberate siderophore-bound iron via siderophore hydrolysis, as shown for FeEnt in *E. coli* (*8*, *9*).

**Fig. 4.**
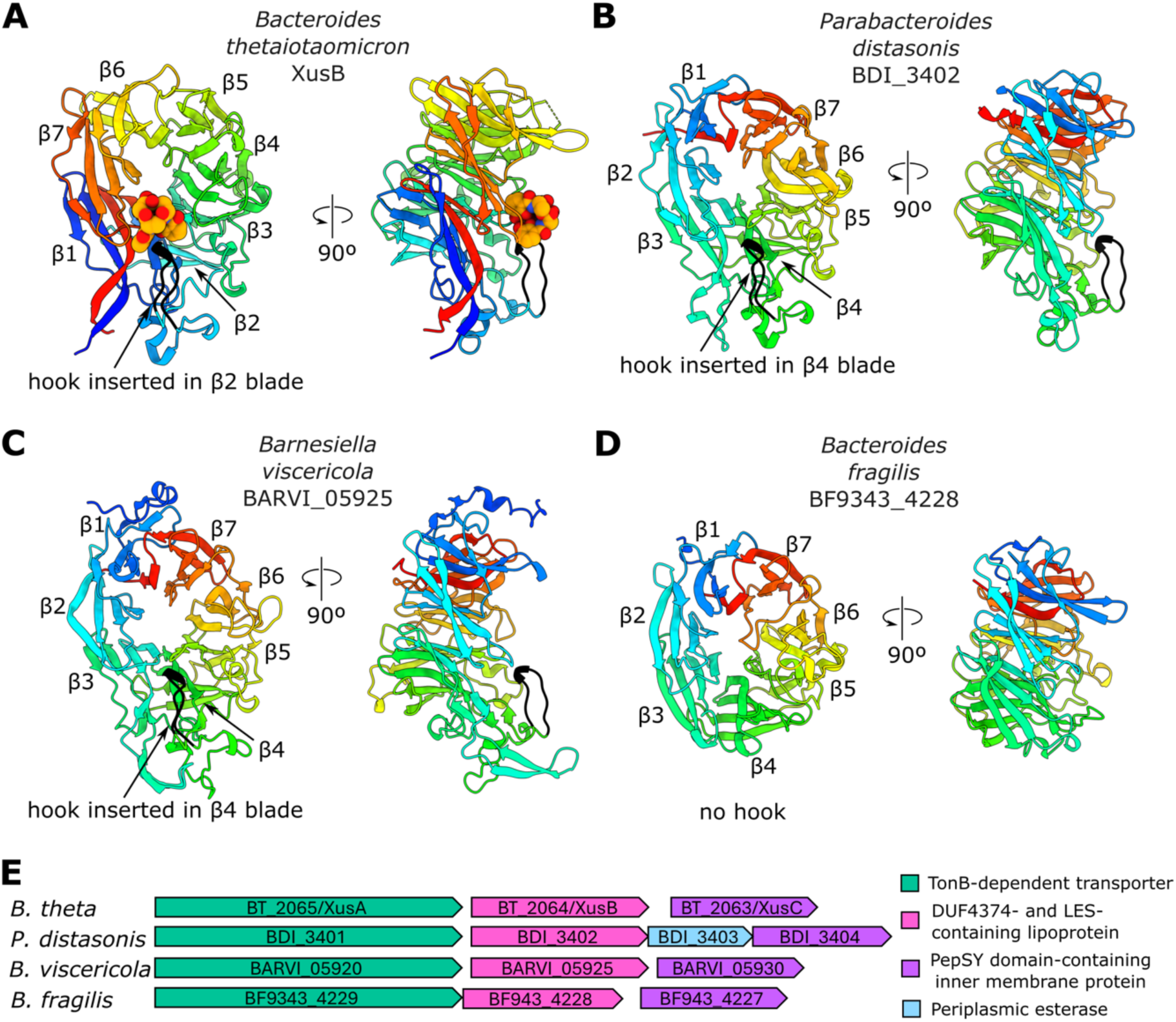
Structural variety of DUF4374-containing proteins. (**A**) BtXusB-FeEnt co-crystal structure. FeEnt atoms are displayed as spheres. (**B**) *P. distasonis* DSM 20701 BDI_3402 (59.52% identity to *B. theta* XusB), (**C**) *B. viscericola* DSM 18177 BARVI_05925 (43.36% identity to BtXusB), and (**D**) *B. fragilis* NCTC 9343 BF9343_4228 (20.21% identity to BtXusB) AlphaFold2 (*36*) models. Views were generated from a superposition, β-propeller blades are labelled β1-7 starting from the N-terminus. All models are coloured in rainbow, from the N-terminus in blue to the C-terminus in red; the hook is in black. AlphaFold2 model prediction confidence is shown in Fig. S8. (**E)** Genetic context of the selected XusB homologues. DUF, domain of unknown function; LES, lipoprotein export signal (*33*).

Conservation of the hook and L2 amino acid sequences between the *B. theta, P. distasonis* and *B. viscericola* homologues (Fig. S7) suggests that, despite differences in position of the hook within the β-propeller, *P. distasonis* and *B. viscericola* homologues might still bind FeEnt, FeDGE and perhaps other catecholate siderophores. However, the *B. fragilis* homologue is unlikely to bind FeEnt due to lack of all FeEnt-binding regions observed in the BtXusB-FeEnt co-crystal structure, which is consistent with a previous report that *B. fragilis* cannot utilize iron bound to catecholate xenosiderophores (*35*).

The amino acid sequence length of BtXusB BLAST hits follows a bimodal distribution with peaks at 410 and 465 amino acids (Fig. S9). *B. theta, P. distasonis* and *B. viscericola* homologues (464, 471 and 491 residues, respectively) belong to the longer group, while the *B. fragilis* homologue (406 residues) belongs to the shorter group. Together with the differences in the xenosiderophore-binding loop regions, the bimodal distribution suggests there are at least two different subtypes of DUF4373-containing proteins.

### *B. viscericola* XusB binds ferric enterobactin

We produced recombinant XusB from *B. viscericola* (BvXusB) to investigate its xenosiderophore-binding properties. We observed binding of FeEnt to BvXusB in ITC with similar affinity to BtXusB (Fig. 5A). We attempted to investigate the molecular details of the interaction between BvXusB and FeEnt and obtained the crystal structure of apo BvXusB (Fig. 5B and Table S1), which confirmed the computational prediction. However, we could not co-crystallise BvXusB with FeEnt, and BvXusB crystals soaked with FeEnt did not diffract. Superposition of the BtXusB-FeEnt and apo BvXusB structures suggests that most BtXusB residues involved in FeEnt binding are present in BvXusB, even though the hook loop in BvXusB is inserted in the β4 blade rather than in the β2 blade as in BtXusB (Fig. 5C). Based on these structural similarities and AlphaFold 3 predictions of BvXusB in complex with FeEnt and FeDGE (Fig. S10), we expect that BvXusB and other XusB homologues with the hook inserted in the β4 blade interact with FeEnt and FeDGE in a very similar manner to BtXusB.

**Fig. 5.**
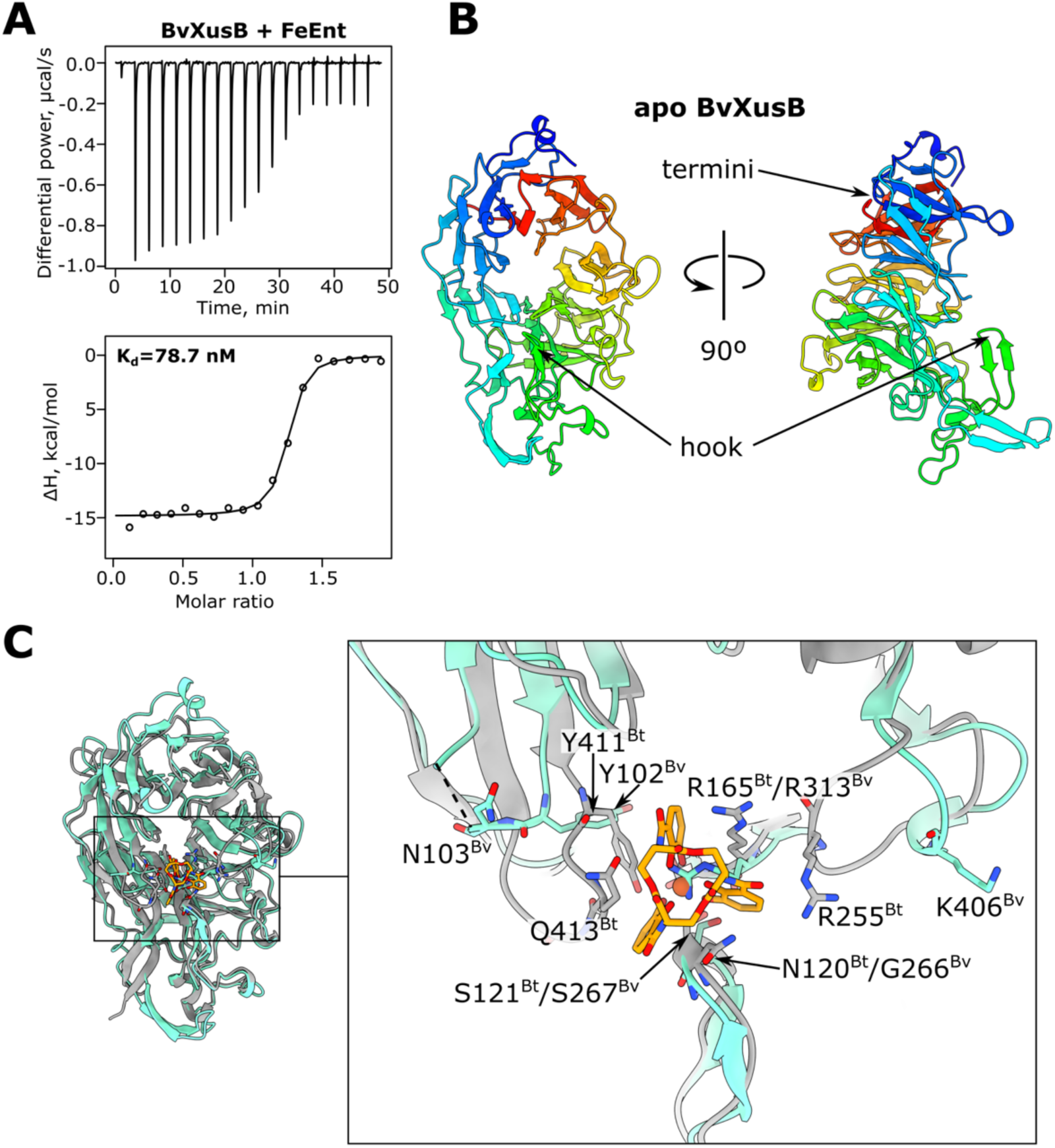
*B. viscericola* XusB binds FeEnt. (**A**) Representative ITC experiment where 250 μM FeEnt was titrated into 25 μM BvXusB (n=2). Integrated heats were fitted to a single binding site model, giving the apparent K_d_ value. (**B**) Crystal structure of apo BvXusB to 3.2 Å. (**C**) Comparison of apo BvXusB (cyan) and BtXusB-FeEnt (grey and orange) crystal structures. The models were superposed using the Matchmaker tool in ChimeraX (Cα-Cα RMSD between 252 pruned atom pairs was 1.0 Å; across all 301 pairs—3.6 Å). BtXusB residues involved in FeEnt binding and the equivalent residues in BvXusB are shown as stick models. The dashed line indicates a disordered loop in the BvXusB crystal structure.

### *B. fragilis* XusB binds ferrichrome

We produced recombinant *B. fragilis* XusB (BfXusB) and determined its crystal structure using data to 1.77 Å, which was consistent with the AlphaFold 2 prediction (Fig. 4D and 6A). BfXusB lacks the hook loop observed in BtXusB crystal structures (Fig. 6B). A previous study reported that *B. fragilis* can grow on ferrichrome as the sole iron source, but the ferrichrome transporter could not be conclusively identified (*35*). Ferrichrome binds iron via hydroxamate groups, rather than catecholate groups as in Ent and DGE (Fig. 6C). FeEnt titrations into BfXusB did not result in substantial injection heats indicating lack of binding, but titrations of ferrichrome into BfXusB showed binding (Fig. 6D,E). Titration of ferrichrome into BtXusB and BvXusB did not show heat changes that would suggest binding (Fig. 6E and Fig. S11). We then determined the structure of BfXusB from crystals soaked with ferrichrome using data to 3.32 Å resolution (Fig. 6F,G and Table S1). We saw additional electron density consistent with the structure of ferrichrome inside the β-propeller pocket without any changes in protein conformation compared to the apo structure (Fig. 6G,H). The moderate resolution makes it difficult to reliably discern which residues of BfXusB interact with ferrichrome, but it is clear the binding residues are completely different to those in BtXusB and BvXusB. The side chains of L93 and W235 slot between two grooves formed by the hydroxamate arms, while the third groove remains unoccupied and exposed to solvent. The side chains of Y52, Y77 and F398 and backbones of W323 and D352-G354 interact with the cyclic peptide backbone (Fig. 6H), in contrast to the BtXusB-FeEnt interaction where there are no contacts with the triserine lactone ring (Fig. 1F). Despite these differences, the xenosiderophore binding sites in BfXusB and BtXusB occupy the same location in the context of the DUF4374 fold (Fig. 6I). This points to a common evolutionary ancestry despite low sequence homology and suggests that other DUF4374-containing proteins also bind their ligands at the same site.

**Fig. 6.**
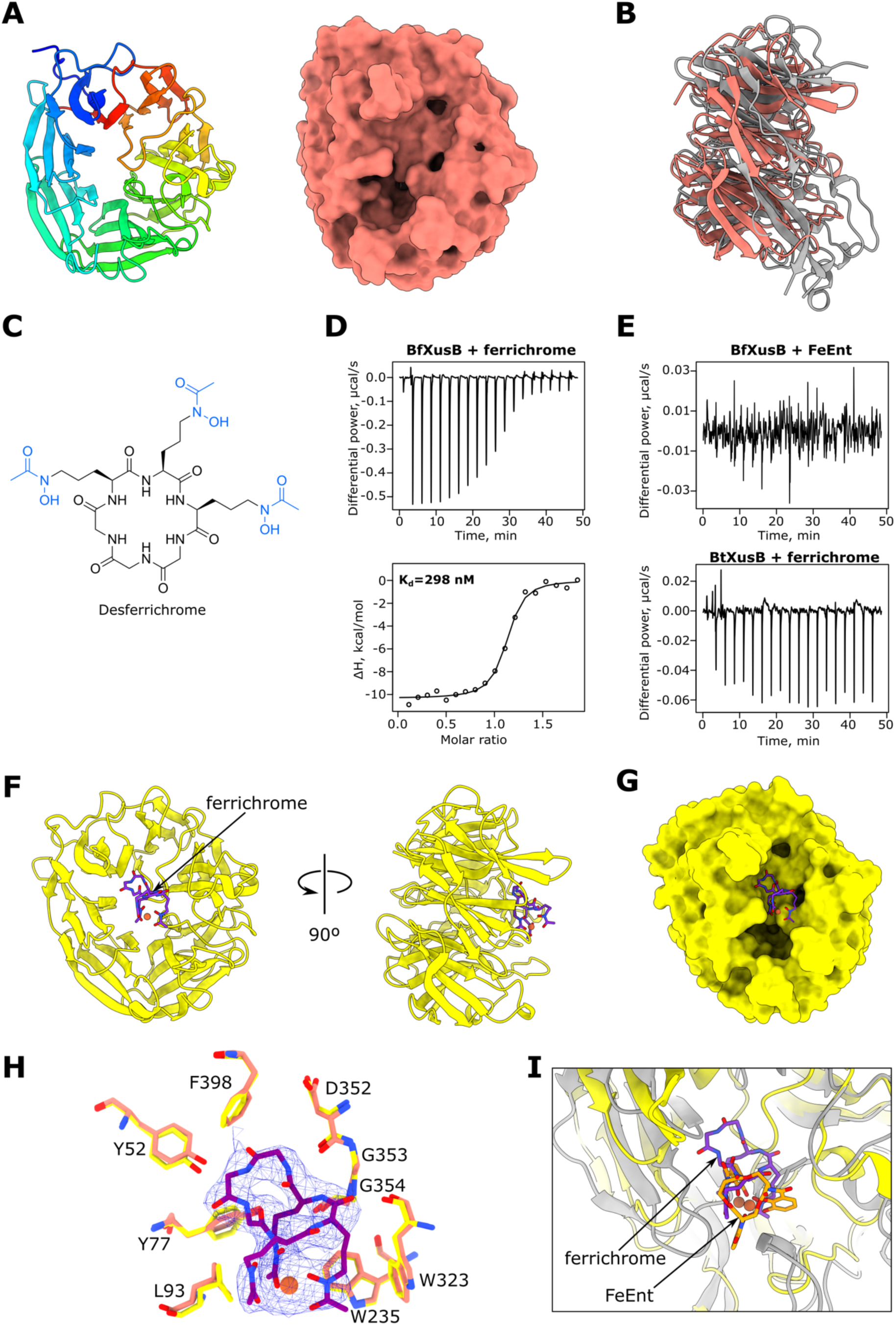
*B. fragilis* XusB binds ferrichrome. (**A**) Crystal structure of apo BfXusB at 1.77 Å shown in cartoon (left) and surface (right) representations. (**B**) Superposition of apo BfXusB and apo BtXusB crystal structures generated using Matchmaker in ChimeraX (Cα-Cα RMSD = 1.2 Å between 140 pruned atom pairs; 3.8 Å across all 249 pairs). (**C**) Chemical structure of desferrichrome. The ferric iron-ligating hydroxamate groups are coloured in blue. (**D**) Representative ITC experiment where 250 μM ferrichrome was titrated into 25 μM BfXusB (n=4 repeats). Integrated heats were fitted to a single binding site model, giving the apparent K_d_ value. (**E**) Representative ITC experiments where 250 μM FeEnt was titrated into 25 μM BfXusB (n=2 experiments) and 250 μM ferrichrome was titrated into 25 μM BtXusB (n=2 experiments). (**F**) Crystal structure of ferrichrome-bound BfXusB to 3.32 Å. (**G**) Surface representation of BfXusB with ferrichrome shown as a stick model inside the shallow β-propeller pocket. (**H**) Ferrichrome model fit into the 2mF_o_-DF_c_ electron density map at 1σ. Residues that likely interact with ferrichrome are shown as yellow stick models. The same residues in the apo BfXusB crystal structure are shown as salmon stick models. (**i**) Superposition of BfXusB-ferrichrome (yellow and purple) and BtXusB-FeEnt (grey and orange) crystal structures (Cα-Cα RMSD between 140 pruned atom pairs was 1.2 Å; across all 249 pairs—3.8 Å). The distance between the two Fe^3+^ ions in the aligned structures is 2.3 Å.

### Xenosiderophore utilization bioassay

The BfXusB-ferrichrome structure and binding data strongly suggested that *B. fragilis* takes up ferrichrome across its outer membrane via a XusAB-like complex formed by BfXusB and a XusA homologue encoded in the same operon (BF9343_4229; Fig. 4E). We wanted to experimentally confirm this and used a modified assay by Rocha & Krykunivsky (*35*) to test xenosiderophore utilization in *B. theta* and *B. fragilis xusAB* deletion strains. We made *xusAB* knockouts in *B. theta* and *B. fragilis* thymidine kinase deletion (*tdk^-^*) strains using the pExchange-tdk allelic exchange system (*37*). As shown previously (*21*), *B. theta* could grow under iron-limiting conditions on both FeEnt and FeDGE as the sole iron source in a *xusAB-* dependent manner (Fig. 7A). We found that growth of *B. fragilis* on ferrichrome is also *xusAB*-dependent (Fig. 7B). This observation confirms the in vivo relevance of our in vitro findings. Surprisingly, we found that *B. fragilis* with and without the *xusAB* locus could utilize FeEnt (Fig. 7A). This result implies that there are more xenosiderophore utilization systems in *B. fragilis*, though BLAST searches did not reveal any paralogous loci to *xusABC* (BF9343_4229-4227).

**Fig. 7.**
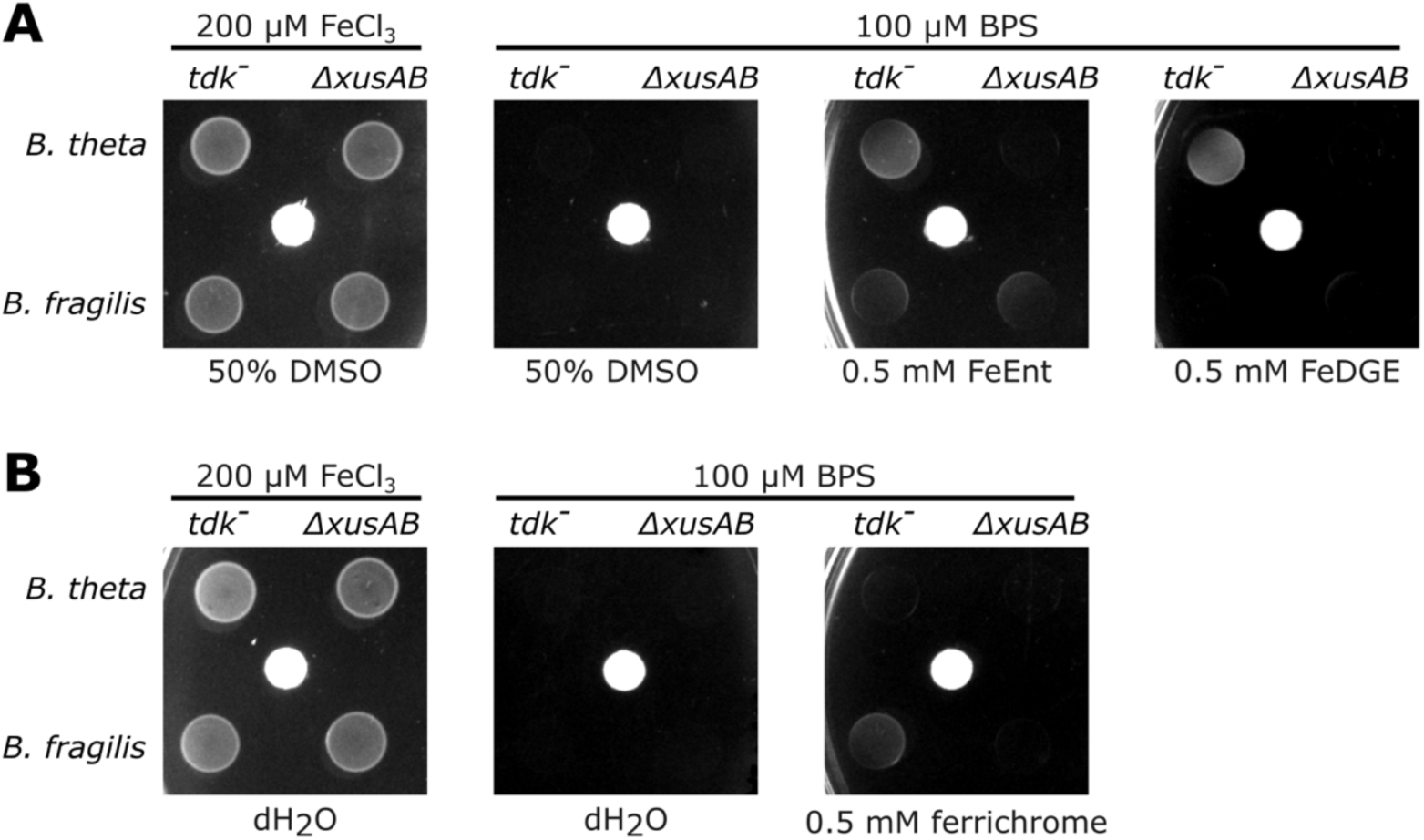
Xenosiderophore utilization bioassay. *B. theta* and *B. fragilis tdk^-^* and Δ*xusAB* strains were spotted on minimal medium agar plates under iron-replete (200 μM FeCl_3_) or iron-limiting conditions (100 μM BPS). Iron-siderophores or the solvent controls were supplied via sterile filter discs (white). The *tdk^-^* strains are thymidine kinase knockouts that are sensitive to 5-fluoro-2′-deoxyuridine that is used for counter-selection during allelic exchange but otherwise have wild-type genotypes. (**A)** shows FeEnt and FeDGE (both supplied in 50% DMSO) utilization. (**B**) shows ferrichrome utilization.

## Discussion

There are many TonB-dependent transporters in Bacteroidota that transport unknown molecules (*38*). It is important to elucidate what these transporters are transporting and how, so that we can obtain a more complete understanding of microbial interactions inside the gut. Our study provides insights into xenosiderophore uptake via Bacteroidota TonB-dependent transporter-lipoprotein complexes and contributes to the ever-widening substrate range described for lipoprotein-assisted TonB-dependent transporters (*2*, *28*, *29*, *38–41*).

We observed an excess of XusB in our native XusAB purifications (Fig. S3). The excess XusB is likely destined for secretion in outer membrane vesicles (OMVs) (*22*). It has been shown that OMVs containing FeEnt-loaded XusB can be used as an iron source by *B. theta* (*22*). This observation implies that the OMVs can somehow deliver FeEnt bound to XusB to XusA in the OM of recipient bacteria. However, our cryo-EM structure shows that XusAB form a tight, closed complex even in the absence of FeEnt. It is unclear how the FeEnt bound to XusB could access the apo-XusB bound to XusA. We speculate that XusB transiently opens like a lid and gives access to the XusAB cavity to incoming xenosiderophores, analogous to the mechanism proposed for vitamin B_12_ import by *Bacteroides* (*28*). Alternatively, the disordered β9-10 extracellular loop of XusA might act as a lateral gate for ingress of xenosiderophores. The latter hypothesis is favoured by lack of any XusAB particles in an open state in the cryo-EM data and the many interactions between the closed XusB lid and XusA (Fig. S5). On the other hand, closure of a mobile XusB lid appears more effective in dislodging the XusB-bound siderophore via clashes with XusA loop β7-8, and we therefore favour the hypothesis of a mobile XusB lid delivering the xenosiderophore to the XusA transporter. It is noteworthy that, while open lipoprotein lids have been observed experimentally for all characterised dimeric SusCD transporters (*29*, *39–41*), for unknown reasons this has not been the case for BtuBG (*28*) and XusAB systems, which are all monomeric. It is also unclear why we could not load the purified XusAB complex with FeEnt or FeDGE. The in vitro conditions may somehow preferentially stabilise the detergent-purified complex in a closed state, while in vivo the presence of negatively charged lipo-oligosaccharide in the *B. theta* OM could destabilise the XusAB interaction. It is also conceivable that excess free, but OM-anchored XusB plays a role in complex opening.

*B. theta* XusAB can transport at least two structurally similar xenosiderophores, FeEnt and FeDGE (*21*, *35*). BtXusB interacts extensively with the siderophore triscatecholate structure, and the binding site can also accommodate the two glucosyl groups of FeDGE. Furthermore, the triserine lactone of FeEnt and FeDGE is exposed to solvent, suggesting that the binding site could accommodate siderophores with different backbones connecting the catecholate arms, such as vibriobactin and bacillibactin, or synthetic Ent mimics (*20*, *42*). This interaction mode raises the question of how many different xenosiderophores a single XusAB complex can import, which could have a substantial impact on our understanding of competition for iron at the host-pathogen-commensal interface.

Together, our structures, functional data and bioinformatics analyses suggest that the presence or absence of the hook in DUF4374-containing proteins indicates preference for catecholate or hydroxamate xenosiderophores, respectively. Whether the shorter, BfXusB-like homologues can bind multiple related xenosiderophores as well remains unknown. Cyclic hydroxamate siderophores structurally related to ferrichrome, such as ferricrocin and ferrichrysin, have hydroxymethyl modifications on the peptide backbone (*43–45*). Because BfXusB interacts extensively with the backbone, these substitutions may either form additional contacts with BfXusB or clash with its sidechains. Further functional and structural analyses are required to determine the substrate range of hydroxamate-binding XusB homologues.

While catecholate siderophores are produced by most Enterobacteriaceae, hydroxamate siderophores are mainly produced by fungi (*45*). Our results suggest that many Bacteroidota species, such as *B. fragilis*, can steal fungal siderophores. Interactions between bacteria and fungi in microbial communities are complex. *Candida albicans* has been shown to promote growth of *B. fragilis* in vitro (*46*). On the other hand, bacteria that have type VI secretion systems, including *B. fragilis,* can directly inject antifungal effectors into fungal cells (*47–49*). The role of iron piracy by *B. fragilis* in the context of its interactions with fungi remains unknown and requires further study.

## Methods

### Bacterial strains and culture conditions

All strains used in this study are listed in Table S4. *E. coli* BL21(DE3) (*50*) and TOP10 strains were genetically manipulated using standard laboratory procedures. The kanamycin concentration used in liquid and solid lysogeny broth (LB) was 50 μg/ml. *B. theta* VPI-5482 *tdk^-^* was cultured either in brain-heart infusion (BHI, Oxoid) supplemented with 1 μg/ml hemin or defined minimal medium (50 mM potassium phosphate buffer pH 7.2, 15 mM sodium chloride, 7.5 mM ammonium sulfate, 9.4 mM sodium carbonate, 4.1 mM L-cysteine, 0.37 nM vitamin B_12_, 5.8 µM vitamin K, 180 µM calcium chloride, 100 µM magnesium chloride, 50 µM manganese(II) chloride and 42 µM cobalt(II) chloride)(*51*) supplemented with 1 μg/ml hemin and 0.5% fructose*. B. theta* was grown under anaerobic conditions at 37 °C in a Don Whitley A35 workstation. When required, 200 µg/ml gentamycin and 25 µg/ml erythromycin was used for selection of *B. theta*.

### Construction of expression plasmids

All plasmids used in this study are listed in Table S5. The nucleotide sequence coding for amino acid residues 35-464 of *B. theta* VPI-5482 XusB (*bt_2064*) and 30-406 of *B. fragilis* NCTC 9343 XusB (*bf9349_4228*) which excludes the signal sequence, the lipid anchor cysteine and a disordered linker, was amplified by PCR using genomic DNA as the template and primers that introduce overhangs containing NcoI and XhoI restriction sites. The PCR product was digested with NcoI and XhoI FastDigest restriction enzymes (ThermoFisher Scientific) and ligated into pET28b, resulting in a C-terminal His_6_-tag fusion. The XusB^R167A,Δhook^ binding site variant nucleotide sequence with flanking NcoI and XhoI restriction sites was synthesised and ligated into the pTwist Amp High Copy plasmid by Twist Bioscience. The synthetic DNA sequence was sub-cloned into pET28b using the flanking restriction sites. Similarly, the nucleotide sequence encoding amino acid residues 29-491 of *B. viscericola* DSM 18177 XusB (*BARVI_05925*) was synthesised by Twist Bioscience and sub-cloned into pET28b using NcoI and XhoI restriction sites. Cloning was carried out in *E. coli* TOP10 cells. Clones were screened for successful insert ligation by colony PCR using EmeraldAmp GT PCR master mix (Takara Bio) with T7 promoter and T7 terminator primers. All constructs were verified by Sanger sequencing (Eurofins).

### Construction of chromosomal deletions

*B. theta* and *B. fragilis xusAB* chromosomal deletions were generated using the pExchange-tdk plasmid (*37*). The pExchange-tdk plasmids containing ∼700 bp flanking the *xusAB* coding sequences in *B. theta* VPI-5482 (*bt2064-65*) or *B. fragilis* NCTC 9343 (BF9343_4228-29) were used to transform *E. coli* S-17 λ pir cells (*52*). The *E. coli* cells were used to introduce the pExchange-tdk plasmids into recipient *Bacteroides* strains lacking thymidine kinase (*tdk^-^*). Conjugants, which underwent a single recombination event, were selected on BHI-hemin plates containing gentamicin (200 μg/ml) and erythromycin (25 μg/ml). Single colonies were cultured in enriched BHI medium overnight, pooled, and plated on BHI-hemin agar plates containing 5-fluoro-2′-deoxyuridine (FUdR; 200 μg/ml) to select for cells that have eliminated the vector backbone from their genome in a second recombination event. After 48 h of growth, FUdR-resistant colonies were restreaked on fresh BHI-hemin-FUdR plates. Clones were screened for successful *xusAB* deletion using PCR with primers that bind outside the homologous region used to make the pExchange-tdk plasmids and confirmed by Sanger sequencing (Eurofins).

### Protein expression and purification in *E. coli*

*E. coli* BL21(DE3) cells were transformed with plasmids carrying the coding sequences for soluble variants of XusB, plated on LB kanamycin agar plates and incubated at 37 °C overnight. The following day, approximately one third of transformants were scraped off the agar plate and used to start an LB kanamycin preculture, which was incubated at 37 °C with shaking for 1.5-3 hours. 12-15 ml of preculture was used to inoculate 1 l flasks of pre-warmed LB kanamycin medium. The cultures were incubated at 37 °C with shaking until OD_600_ reached 0.4-0.8. Protein expression was then induced by adding isopropyl β-d-1-thiogalactopyranoside to a final concentration of 0.1 mM, and the cultures were grown for another 16-20 h at 18 °C with 150 rpm shaking. Cells were harvested by centrifugation at 8,000 × *g* at 4 °C for 20 min. Pellets were resuspended in cold Tris-buffered saline (TBS, 20 mM Tris-HCl, 300 mM NaCl) and stored at -20 °C until protein purification. Cell pellets were thawed and homogenized in TBS using a Dounce tissue grinder and supplemented with DNase I (Roche). Cells were lysed by passing the suspension once through a cell disruptor (Constant Systems) at 20 kpsi. The lysates were supplemented with 1 mM phenylmethylsulfonyl fluoride (PMSF) and clarified by centrifugation at 30,000 *g* for 30 min at 4 °C. The supernatants were loaded on 5 ml chelating Sepharose columns charged with Ni^2+^ ions using gravity flow. The columns were washed with 30 column volumes of TBS with 30 mM imidazole. Bound proteins were eluted with 4 column volumes of TBS with 250 mM imidazole. The eluate was concentrated with an Amicon Ultra filtration device (30 kDa cut-off membrane), loaded on a HiLoad Superdex 200 16/600 pg column and eluted in 10 mM HEPES-NaOH pH 7.5, 100 mM NaCl. Fractions were analysed by SDS-PAGE. Fractions containing the soluble XusB variants were pooled, concentrated, flash-frozen and stored at -80 °C.

### Synthesis of DGE

DGE was prepared chemoenzymatically based on previously reported procedures (*53*, *54*). A reaction solution containing 500 μM Ent and 3 mM UDP-glucose (UDP-Glc) in buffer (75 mM Tris-HCl pH 8.0, 5 mM MgCl_2_, 2.5 mM TCEP) was prepared and aliquoted into 10 × 750 μl portions. The *C-*glucosyltransferase IroB was added to each sample to a final concentration of 5 μM and the reactions were incubated at room temperature. After 1 h, each aliquot was quenched by addition of 75 μl 6% trifluoroacetic acid (TFA)/H_2_O and diluted with ∼300 μl acetonitrile (MeCN). Quenched solutions were centrifuged (13,000 rpm, 10 min, 4 °C) then purified by preparative reversed phase HPLC (0−100% B in 30 min, H_2_O/MeCN + 0.1% TFA, 10 mL/min). The peak corresponding to DGE was collected and lyophilized to dryness.

### Preparation of ferric siderophores

Ent and DGE were dissolved in dimethyl sulfoxide (DMSO) to make 10 mM solutions. The 10 mM siderophore solution was mixed in a 1:1 volume ratio with 10 mM aqueous FeCl_3_ solution to yield a 5 mM ferric siderophore stock solution in 50% (v/v) DMSO. The concentrations of FeEnt and FeDGE stock solutions were determined spectrophotometrically using the extinction coefficient ε_495_ = 5,600 M^−1^ cm^−1^ (in 20 mM Tris-HCl pH 7.0, 50% methanol)(*55*). The stock solutions were stored in small aliquots at -80 °C to minimise siderophore hydrolysis. Similarly, desferrichrome (Merck) was dissolved in sterile Milli-Q water to make a 10 mM solution, aliquoted and stored at -20 °C. After thawing, 10 mM desferrichrome aliquots were mixed with 10 mM FeCl_3_ in a 1:1 ratio, resulting in a 5 mM aqueous ferrichrome solution. Ferrichrome concentration was confirmed spectrophotometrically using the extinction coefficient ε_425_ = 2,900 M^−1^ cm^−1^ (in 20 mM Tris-HCl pH 7.0)(*56*).

### Crystal structure determination

The purified soluble BtXusB variant was concentrated to 36 mg/ml. Sitting drop vapour diffusion crystallisation screens were set up using a Mosquito robot (SPT Labtech) either with apo BtXusB or with a 1:1.1 molar ratio of BtXusB and FeEnt. The crystallization plates were incubated at 20 °C. Apo XusB crystals appeared after less than a week in the PACT premier screen (Molecular Dimensions) containing 0.1 M MES pH 6.0, 0.2 M calcium chloride and 20% PEG 6000. Crystals were cryo-protected in mother liquor supplemented with ∼20% PEG 400 and flash-cooled in liquid nitrogen. BtXusB-FeEnt co-crystals appeared in the Index screen (Hampton Research) condition containing 0.1 M citric acid pH 3.5 and 2 M ammonium sulphate after 2 weeks. Crystals were cryo-protected by passing through a drop of 3.5 M ammonium sulphate and flash-cooled in liquid nitrogen. Apo BtXusB crystals grown in hanging vapour diffusion drops (0.1 M HEPES pH 6.7, 0.2 M calcium chloride, 20% PEG 6000, and 5% DMSO) were soaked with 1 mM FeDGE in mother liquor for 24 h, cryoprotected with mother liquor, 1 mM FeDGE and 20% PEG 400, and flash-cooled in liquid nitrogen.

BvXusB was concentrated to 35 mg/ml and sitting drop crystallization trials using commercial screens were set up as above. An initial hit was observed in the Structure 1+2 screen (Molecular Dimensions) and further optimized using hanging drop vapour diffusion and streak seeding. The final condition for diffracting apo BvXusB crystals was 0.1 M sodium acetate pH 4.8, 0.2 M ammonium sulfate and 35% PEG 2000 monomethyl ether. The crystals were cryoprotected in mother liquor with 20% PEG 400 and flash-cooled in liquid nitrogen.

BfXusB was concentrated to 30 mg/ml and sitting drop crystallization trials using commercial screens were set up as above. Apo crystals were harvested directly from the Index screen (Hampton Research) condition containing 0.1 M HEPES pH 7.5, 0.02 M magnesium chloride hexahydrate, 22% w/v poly(acrylic acid sodium salt) 5100. Mother liquor with 20% PEG 400 was used to cryoprotect the crystals before flash-cooling in liquid nitrogen. Crystals from an identical condition in the JCSG+ screen (Molecular Dimensions) were soaked with 1 mM ferrichrome in mother liquor for 3 weeks, cryoprotected in mother liquor, 1 mM ferrichrome and 20% PEG 400 and flash-cooled in liquid nitrogen.

X-ray diffraction data were collected at the Diamond Light Source synchrotron (UK) at a temperature of -173 °C (Table S1). Datasets were processed with XIA2-dials (*57*), scaled with Aimless (*58*), and the space group was confirmed with Pointless (*59*). The apo structures were solved by molecular replacement with computational models generated by AlphaFold2 (*36*). The siderophore-bound structures were solved by molecular replacement using the apo experimental structures. All models underwent cycles of manual building in Coot (*60*) and refinement in Phenix (*61*) until no further improvement in R factors could be achieved. The models were validated using MolProbity (*62*). Refinement statistics are summarised in Table S1.

### Isothermal titration calorimetry

ITC was carried out in 10 mM HEPES–NaOH pH 7.5 and 100 mM NaCl supplemented with 5% DMSO to improve FeEnt and FeDGE solubility. A 250 μM FeEnt solution was made in ITC buffer, and any precipitate was removed by centrifugation. The clarified FeEnt solution was titrated into 25 μM protein at 25 °C using a Microcal PEAQ-ITC instrument (Malvern Panalytical). After an initial delay of 60 s, a single injection of 0.4 μl was carried out, which was discarded from data analysis, followed by 18 injections of 2 μl spaced in 150 s intervals. For FeDGE titrations only, the initial 0.4 μl injection was followed by 12 injections of 3 μl spaced in 150 s intervals. The sample cell was stirred at 750 rpm during titration. Ligand to buffer control titrations were subtracted from all experiments. The experiments were repeated at least twice (Table S2). Data were fitted to a single-binding-site model using the Microcal PEAQ-ITC Analysis software v1.40. For FeDGE titrations only, the stoichiometry (n) was fixed to one and the ligand concentration was allowed to float as the fits would not converge otherwise. We speculate that the FeDGE concentration may have been inaccurate due to hydrolysis of the siderophore during repeated freeze-thawing. Data fitting results for successful binding experiments are shown in Table S2.

### Growth curve experiments

*B. theta tdk^-^* strain was cultured anaerobically in BHI supplemented with hemin at 37 °C overnight. 0.2 ml of the overnight culture was used to inoculate 5 ml of fresh supplemented BHI the next morning, followed by a 4 h incubation. The cells were collected by centrifugation for 5 min at 2,800 × *g*, 20 °C, and resuspended in 1 ml of fresh, pre-warmed minimal medium. Minimal medium supplemented with fructose and hemin was aliquoted into tubes, to which BPS, FeEnt and FeCl_3_ was added as required. Washed cells were diluted in the appropriate tubes to OD_600_=0.04 and dispensed in 200 μl aliquots into the wells of a sterile 96-well plate in triplicate. Growth at 37 °C was monitored for 48 h using a Biotek Epoch microplate reader housed inside an anaerobic workstation. The experiment was repeated three times with similar results.

### Purification of the XusAB complex from *B. theta*

A C-terminal His_6_-tag was fused to XusB by introducing the tag coding sequence into the *B. theta tdk^-^* chromosome before the stop codon of *bt_2064* via allelic exchange using the pExchange plasmid (*37*). Presence of the tag on the chromosome was confirmed by PCR and Sanger sequencing. Conditions under which the tagged XusB is expressed were identified by Western blotting. The *B. theta bt_2064-his* strain was cultured anaerobically in tubes containing 2 ml minimal medium supplemented with 6.25-100 μM BPS for 18 h. Equivalents of 1 ml culture at OD_600_=2 were pelleted by centrifugation. The pellets were resuspended in 80 μl BugBuster (Sigma), supplemented with 1 mM PMSF, and incubated for 15 min at room temperature. Cell debris was pelleted by centrifugation in a benchtop microcentrifuge. Samples from cells grown in the presence of different amounts of BPS were separated by SDS-PAGE and transferred onto a PVDF membrane via wet transfer. The PVDF membrane was stained with Ponceau S stain to confirm successful transfer and blocked with 1% milk solution in PBS supplemented with 0.1% (v/v) Tween 20 for 20 min at room temperature. The membrane was probed with anti-His-horseradish peroxidase conjugate antibody (Roche; 1:500 dilution in 1% milk solution) for 1 h at room temperature and washed three times with PBS-Tween. The blots were developed using SuperSignal West Pico Plus chemiluminescent substrate (Thermo Fisher Scientific) and imaged using a Gel Doc XR+ system (Bio-Rad).

Frozen *B. theta bt_2064-his* glycerol stocks were used to inoculate BHI, followed by overnight incubation at 37 °C under anaerobic conditions. After autoclaving, minimal medium was stored in an A35 Don Whitley anaerobic workstation overnight. The following day, the minimal medium was supplemented 50 μM BPS, fructose (0.5%) and hemin (1 μg/ml). Overnight *B. theta* BHI cultures were pelleted, resuspended in minimal medium to original volume and used to inoculate 0.5 l bottles of minimal medium at a ratio of 1:250. Bacteria were cultured at 37 °C under anaerobic conditions for 18-20 h. Cultures were pelleted by centrifugation at 6,000g for 30 min at 4 °C, resuspended in TBS, and stored at -20 °C.

Pellets were thawed, supplemented with DNase I and homogenised. Cells were lysed by passing the cell suspension once through a cell disruptor at 22 kpsi. The lysate was clarified by centrifugation at 30,000 × g, 4 °C for 30 min. Membranes were isolated from the clarified lysate by ultracentrifugation at 140,000 × g, 4 °C for 50 min, followed by solubilization in 1.5% lauryldimethylamine N-oxide (LDAO) in TBS for 1 h at 4 °C. Insoluble material was pelleted by centrifugation at 44,000 × g, 4 °C for 30 min. The solubilised material was passed through ∼3 ml of chelating Sepharose resin charged with Ni^2+^ ions using gravity flow. The column was washed with 25 column volumes of TBS with 30 mM imidazole and 0.1% dodecyl-β-D-maltopyranoside (DDM, Anatrace), and bound protein was eluted with TBS supplemented with 200 mM imidazole and 0.03% DDM. The eluate was concentrated using an Amicon Ultra filtration device (100 kDa cut-off membrane), loaded on a Superdex 200 10/300 Increase column and eluted in 10 mM HEPES-NaOH pH 7.5, 100 mM NaCl, 0.03% DDM. Fractions corresponding to the XusAB peak were pooled, concentrated, flash-frozen in liquid nitrogen, and stored at −80 °C.

### Cryo-EM structure determination

Pure XusAB complex at 7 mg/ml was incubated with a four-fold molar excess of FeEnt for 45 min at room temperature. 3.5 μl of the complex was then applied to glow-discharged Quantifoil R1.2/1.3 copper 200 mesh holey carbon grids. The grids were immediately blotted for 1-5 s and plunge-frozen in liquid ethane using a Vitrobot Mark IV (ThermoFisher Scientific) device operating at 4 °C and 100% humidity. The grids were initially screened on a 200kV FEI Glacios microscope at the University of York (UK). Data were collected at the Astbury Centre (Leeds, UK) on a FEI Titan Krios microscope operating at 300 kV using a Falcon 4i direct electron detector (ThermoFisher Scientific) operating in counting mode (Table S3). A total of 6,645 movies were recorded in electron event representation (EER) format at 165,000× magnification, corresponding to a pixel size of 0.74 Å.

The cryo-EM workflow is shown in Fig. S4. All data processing was done in cryoSPARC v4.4.1. Movies were motion-corrected using patch motion correction. CTF parameters were fitted using patch CTF correction. Initially, ∼2,000 particles were picked manually and used to make 2D classes for template picking. 1,110,843 picked particles were extracted in 480-pixel boxes (0.74 Å/pixel), Fourier cropped to a box size of 240 pixels (1.48 Å/pixel) and subjected to two rounds of 2D classification. Ab initio models were generated from classes exhibiting protein density and decoy models were generated from classes containing predominately noise. Particles selected after 2D classification were subjected to heterogeneous refinement against four ab initio models, two of which were decoys. A single class from heterogeneous refinement refined to high resolution in non-uniform refinement (*63*). Most particles clustered into a single class in 3D classification (8 classes in total). Particles from this class were re-extracted in 480- pixel boxes (0.74 Å/pixel) and refined using non-uniform refinement (per-particle defocus, tilt and trefoil refinement enabled), followed by local refinement. Per-particle motion correction was performed using reference-based motion correction, followed by local refinement and re-estimation of per-particle defocus, tilt and trefoil parameters. The particle stack was subjected to a final round of 2D classification with the noise model (sigma) annealing turned off to remove any remaining poor-quality particles. Another round of reference-based motion correction was performed, followed by a final round of local refinement that gave a map with a global resolution of 2.7 Å based on the FSC=0.143 criterion. The final stack had 76,979 particles.

The XusAB protein sequences were supplied to ModelAngelo (*64*) for automated model building. The resulting model was iteratively adjusted in Coot (*60*) and ISOLDE (*65*) and refined using Phenix real space refinement (*61*). The model was validated using MolProbity (*62*). Refinement statistics are shown in Table S3.

### Structure analysis and visualisation

Atomic models, electron density maps and cryo-EM maps were analysed in Coot (*60*) and UCSF ChimeraX (*66*). All figures depicting structural data were generated using UCSF ChimeraX. The ISOLDE (*65*) plugin was used to visualise electron density maps.

### Xenosiderophore utilisation bioassay

*B. theta tdk^-^*, *B. theta* Δ*xusAB*, *B. fragilis tdk^-^* and *B. fragilis* Δ*xusAB* strains were grown overnight in enriched BHI (EBHI; 37 g/l Oxoid BHI powder, 5 g/l yeast extract) supplemented with 1 μg/ml hemin. 0.3 ml of the overnight cultures were used to inoculate 5 ml of fresh EBHI-hemin medium. The strains were cultured anaerobically at 37 °C for 3.5 h. Cells were pelleted, resuspended in 5 ml fresh minimal medium with hemin and 0.5% fructose, and diluted to OD∼0.05. 10 μl aliquots were spotted on minimal medium agar plates supplemented with 0.5% fructose, 10 μg/ml protoporphyrin IX and either 200 μM FeCl_3_ or 100 μM BPS. The spots were placed 1.5 cm from the centre of sterile filter discs embedded in agar. Four 5 μl drops of 0.5 mM FeEnt in 50% DMSO, 0.5 mM FeDGE in 50% DMSO, 0.5 mM ferrichrome in sterile water, 50% DMSO only or sterile water only were added to the filter discs. The plates were incubated at 37 °C anaerobically for 18 h before imaging.

## Acknowledgements

The *B. fragilis tdk^-^* strain was a gift from Janet Quinn (Newcastle). We thank Matthew Nodwell (Simon Fraser University) for providing the Ent used in this study. We also thank Diamond Light Source for access to macromolecular crystallography beamlines (proposals mx-24948 and mx-32736). We acknowledge use of the crystallization facilities and the GPU cluster for cryo-EM data processing at the Newcastle University Structural Biology Facility. We acknowledge use of the Glacios microscope for screening cryo-EM grids at the York Structural Biology Laboratory (University of York). We acknowledge use of the Astbury Centre (University of Leeds) Titan Krios microscope.

## Author contributions

A.S. conceived the project, designed experiments, made constructs, purified and crystallized proteins, made cryo-EM grids and collected cryo-EM data, determined crystal and cryo-EM structures, and wrote the manuscript with input from R.N.M, E.M.N. and B.v.d.B. Y.L.S. and H.M. made expression constructs, purified and crystallized proteins, and carried out ITC titrations, supervised by A.S. and B.v.d.B. R.N.M. synthesized DGE, supervised by E.M.N. A.B. collected X-ray diffraction data and provided computational support. B.v.d.B. acquired funding, designed experiments, and collected cryo-EM data.

## Data availability

Electron microscopy volumes of the XusAB complex have been deposited in the Electron Microscopy Data Bank with the accession code EMD-51210, and the atomic coordinates have been deposited in the Protein Data Bank under the accession code 9GBC. Atomic coordinates and the associated crystallographic structure factors have been deposited in the Protein Data Bank under the following accession codes: 9GCY (apo BtXusB), 9GCZ (BtXusB-FeEnt), 9HQ1 (BtXusB-FeDGE), 9GAR (apo BvXusB), 9HQE (apo BfXusB), and 9HQK (BfXusB-ferrichrome). Other data and materials related to the manuscript are available from the corresponding authors upon request.

---

**Fig. S1.**
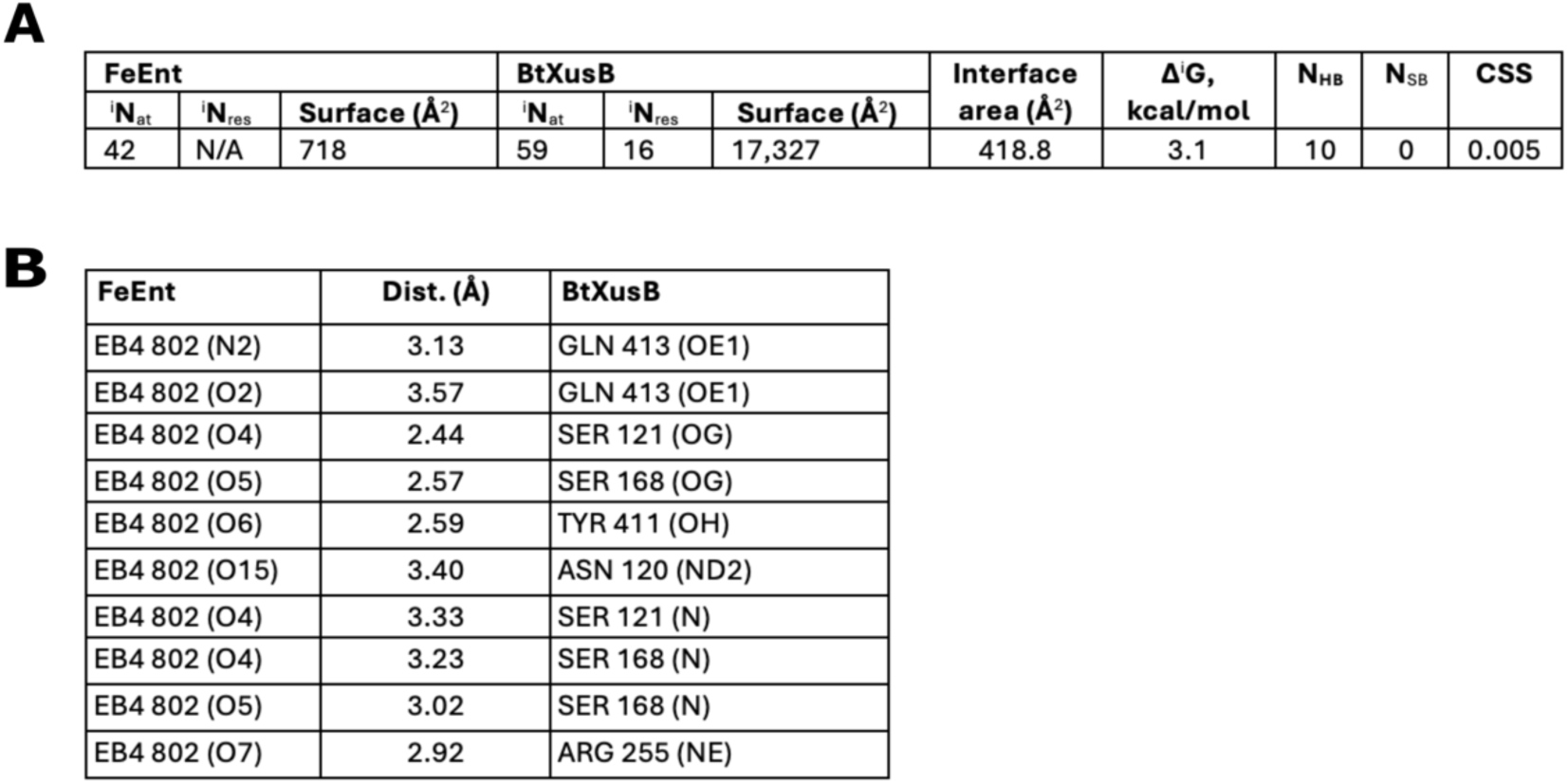
PISA (*1*) analysis of BtXusB-FeEnt interactions. (**A**) Complex surface area calculations. (**B**) Hydrogen bonds formed between FeEnt and BtXusB.

**Fig. S2.**
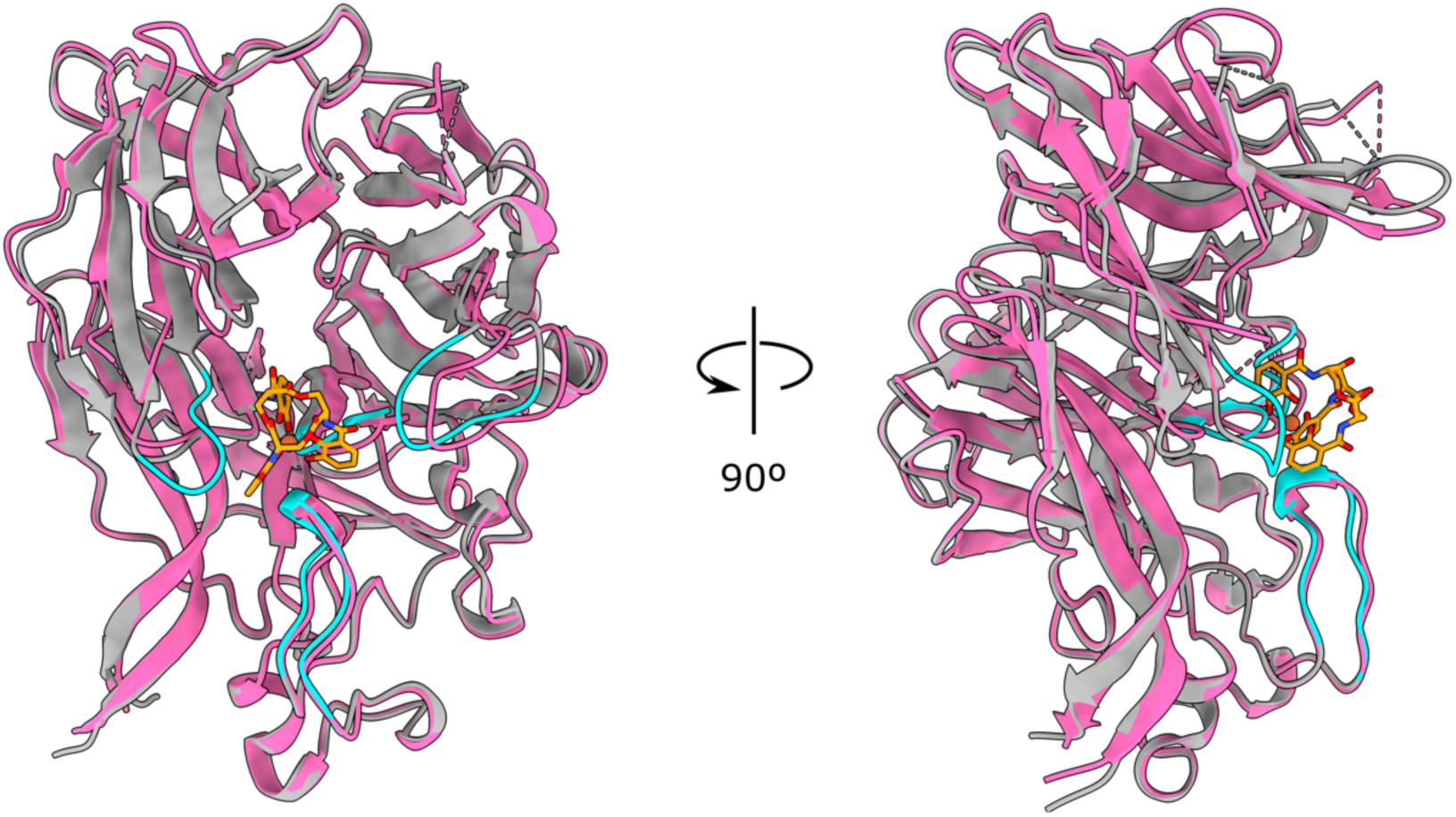
Comparison of apo and FeEnt-bound BtXusB structures. Structural alignment of apo (hot pink) and FeEnt-bound (grey) BtXusB crystal structures. Cα-Cα RMSD = 1.07 Å. FeEnt is in orange; BtXusB FeEnt-binding loops are highlighted in cyan in the co-crystal structure.

**Fig. S3.**
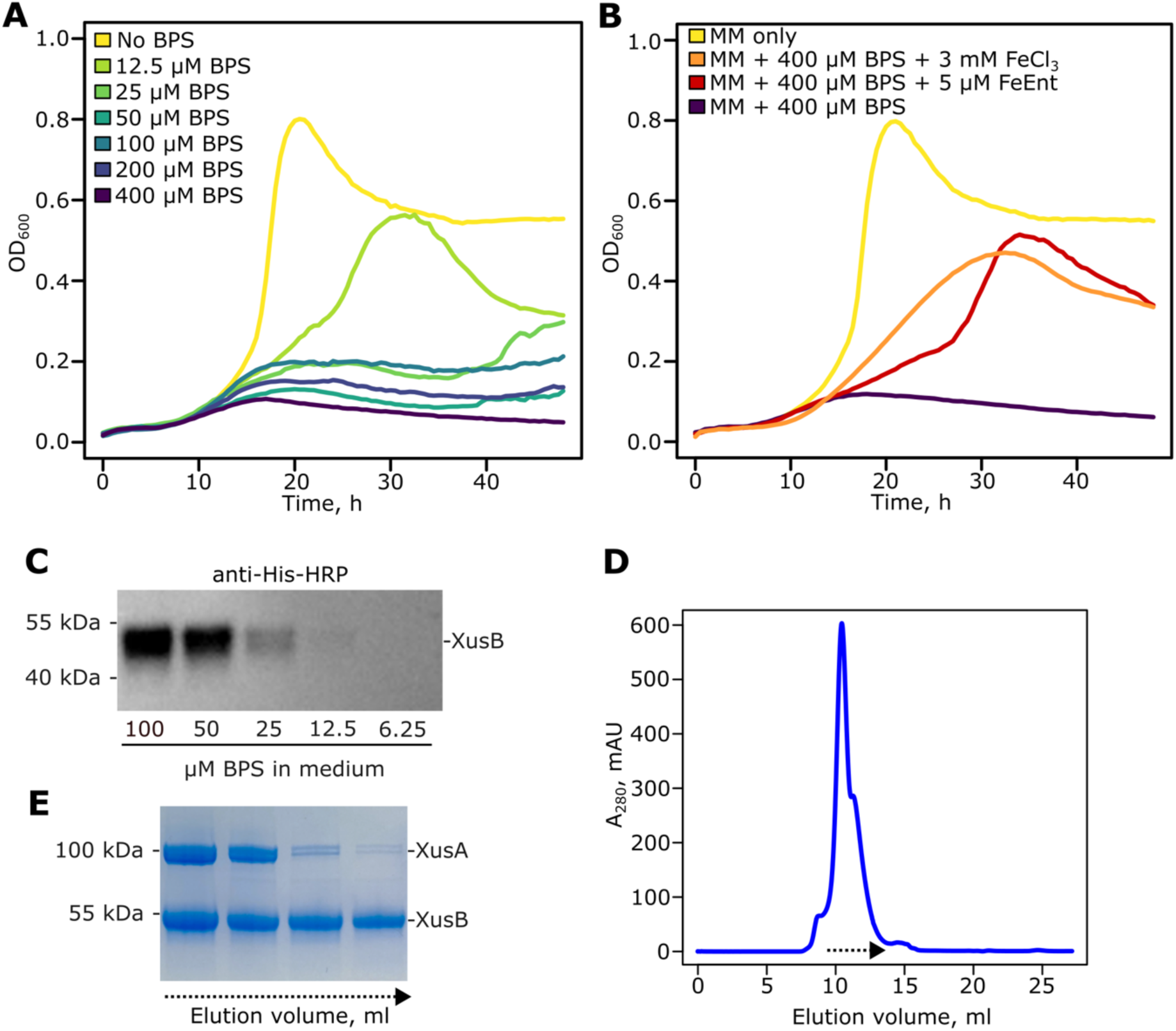
Expression and purification of the native XusAB complex. (**A**) *B. theta tdk^-^* strain grown anaerobically in minimal medium with 0.5% fructose and 1 μg/ml hemin. The iron chelator bathophenanthroline disulfonate (BPS) was added at indicated concentrations. (**B**) *B. theta tdk^-^* strain grown anaerobically in the presence of BPS and indicated iron sources. All growth curves shown are averages from 3 wells of a single 96-well growth experiment. The experiment was repeated three times under identical conditions with similar results. (**C**) Western blot of whole cell lysates from *B. theta bt2064-his* cells grown overnight in minimal medium and the indicated concentrations of BPS. (**D**) Size exclusion chromatography trace of XusAB complex after immobilized metal affinity chromatography on a Superdex 200 10/300 Increase column. (**E**) SDS-PAGE analysis of 0.5 ml elution fractions from (**D**) indicated by the dashed arrow (approximately 10-12 ml elution volume). The indicated bands were identified as XusA and XusB by peptide mass fingerprinting. Both bands around 100 kDa correspond to XusA.

**Fig. S4.**
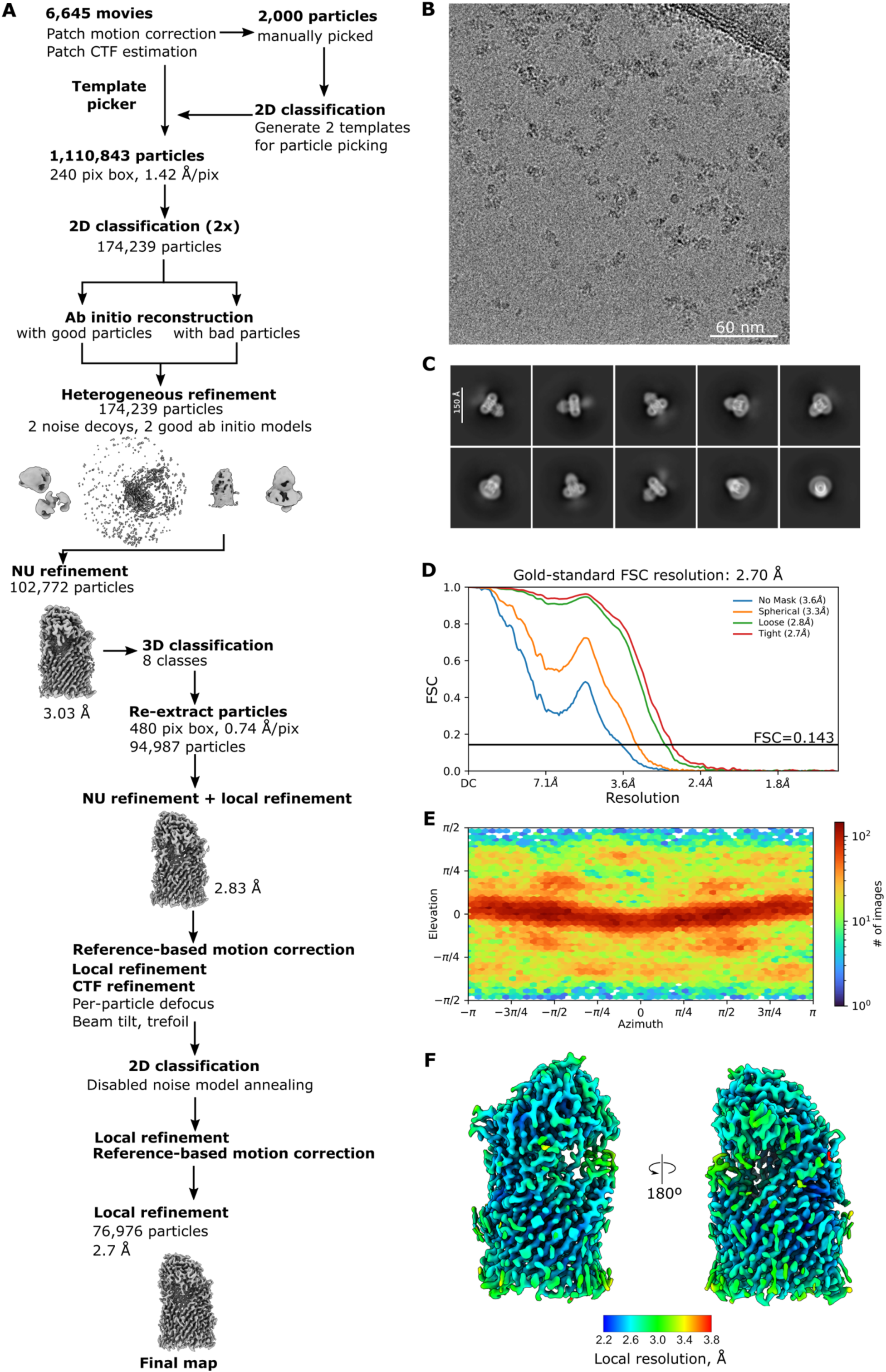
XusAB cryo-EM data processing. (**A**) Data processing workflow carried out in cryoSPARC v4.4.1 (*2*). (**B**) Representative motion-corrected micrograph (n=6,645). (**C**) Representative 2D class averages. (**D**) Gold-standard Fourier shell correlation curve and (**E**) viewing direction distribution plot for the final map. (**F**) The final cryo-EM map coloured by estimated local resolution.

**Fig. S5.**
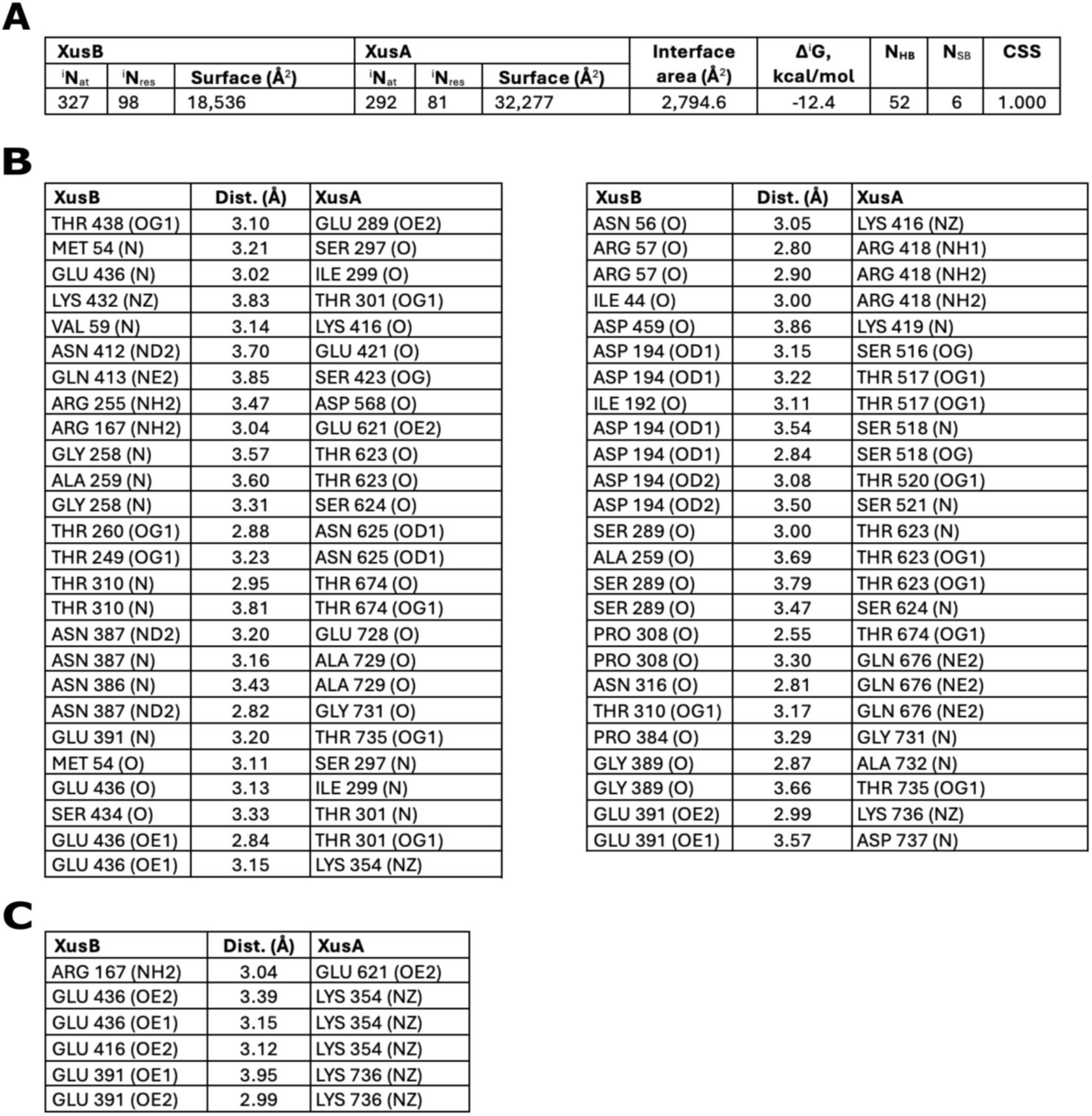
PISA (*1*) analysis of XusA-XusB interactions. (**A**) Complex surface area calculations. (**B**) Hydrogen bonds formed between XusA and XusB. (**C**) Salt bridges formed between XusA and XusB.

**Fig. S6.**
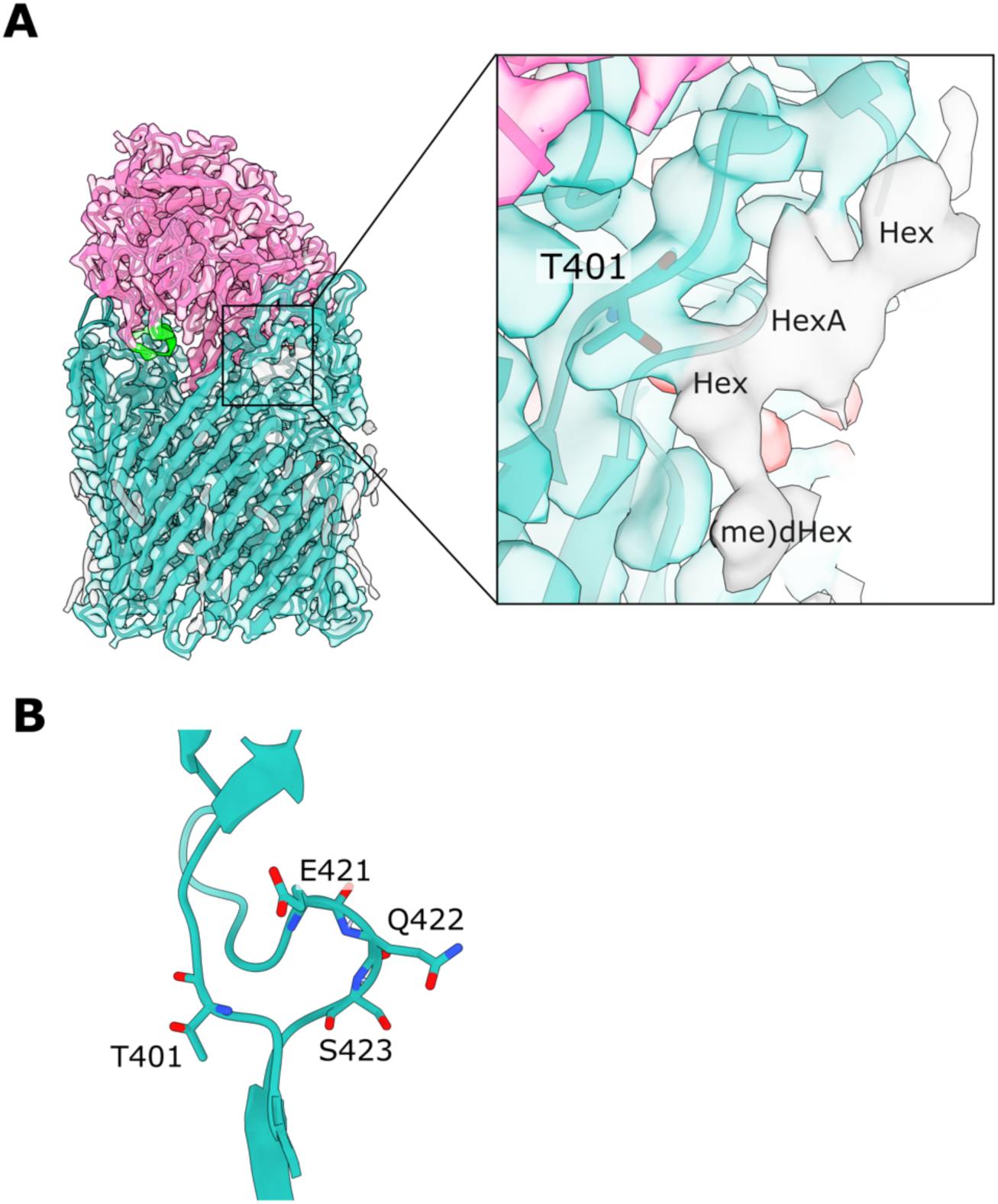
XusA O-glycosylation. (**A**) The glycan density extending from the sidechain hydroxy oxygen of T401 is shown in grey. Sugar units that likely constitute the glycan chain are indicated: Hex, hexose; HexA, hexuronic acid; (me)dHex, (methyl)deoxyhexose. Sugars were assigned based on the structure of the *B. fragilis* O-glycan (*3*). (**B**) Residues 421-423, which are in the C-terminal part of the β7-8 extracellular loop of XusA and are implicated in FeEnt release from XusB, are somewhat close to T401. The sidechain of T401 faces away from residues 421-423 towards the solvent.

**Fig. S7.**
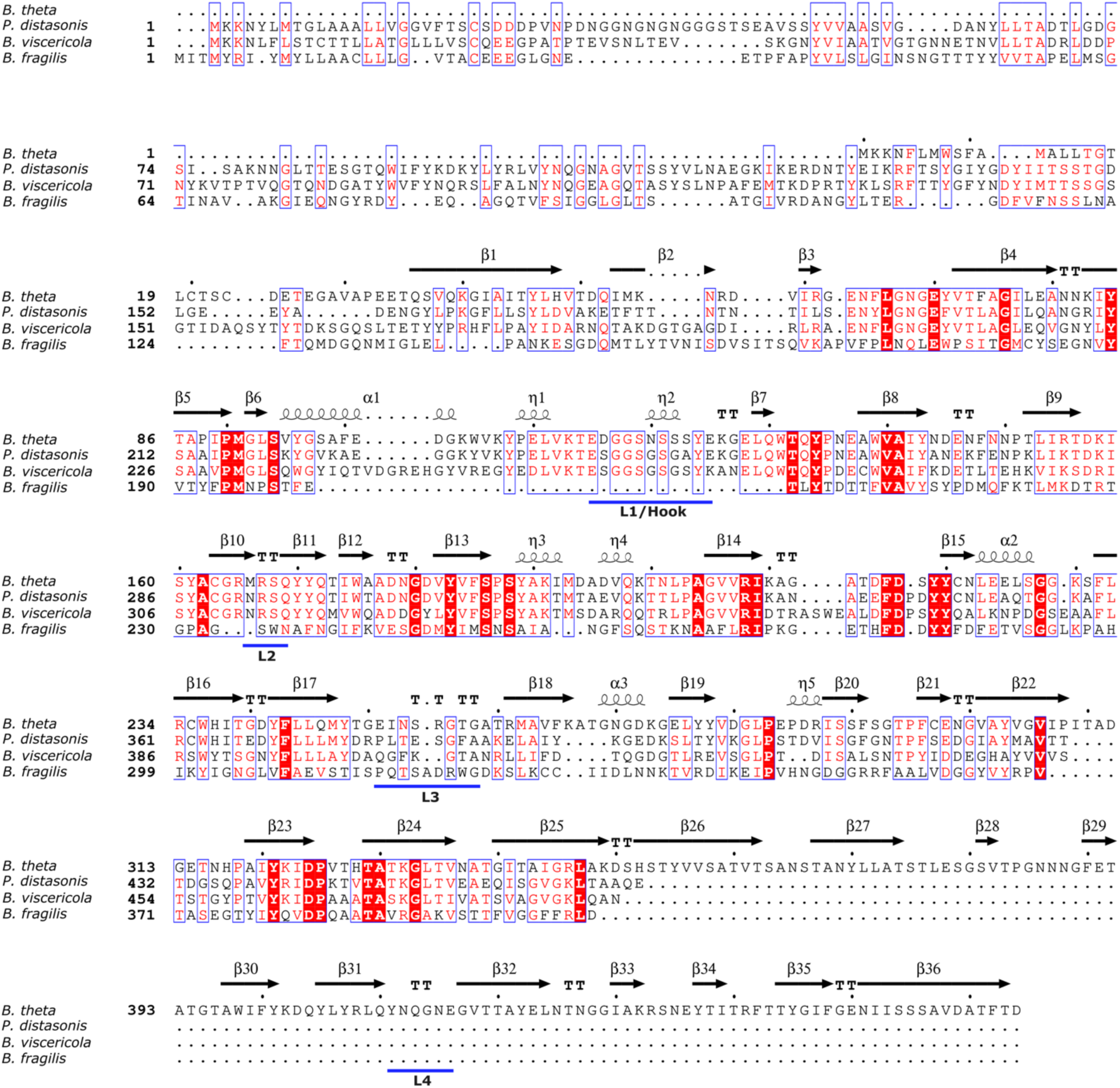
XusB homologue sequence alignment. Alignment of BtXusB, *P. distasonis* DSM 20701 BDI_3402, *B. viscericola* DSM 18177 BARVI_05925, and *B. fragilis* NCTC 9343 BF9343_4228 amino acid sequences. The secondary structure features of BtXusB in the BtXusB-FeEnt structure are annotated above the alignment. Visualised in ESPript v3.0 (*4*) (https://espript.ibcp.fr).

**Fig. S8.**
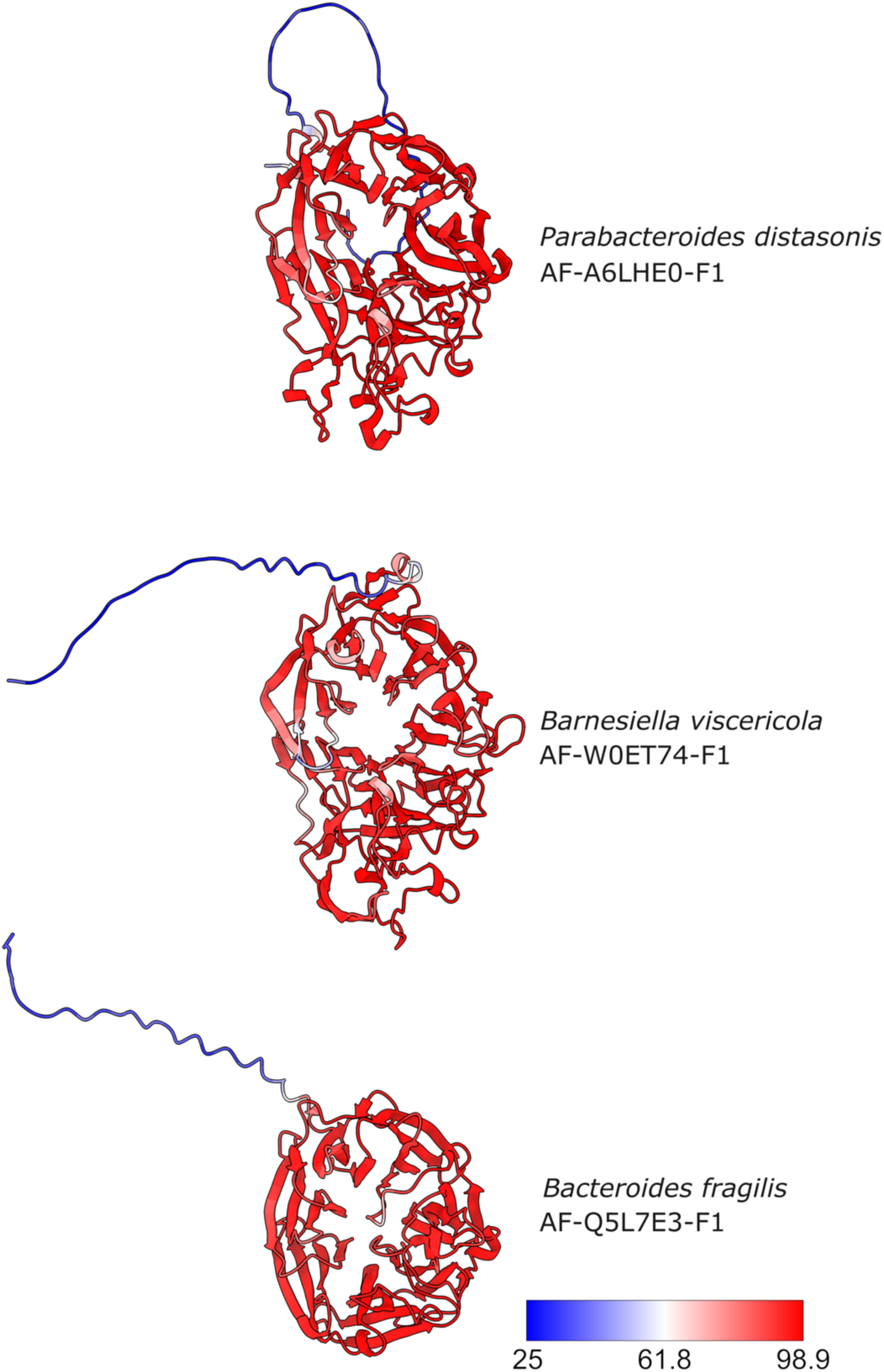
XusB homologue AlphaFold2 (*5*) model prediction confidence. The indicated entries were downloaded from AlphaFold DB. The views were generated from superposition with the BtXusB-FeEnt crystal structure. Each model is coloured according to pLDDT value (colour key). The blue (low confidence) N-terminal regions correspond to the signal peptides.

**Fig. S9.**
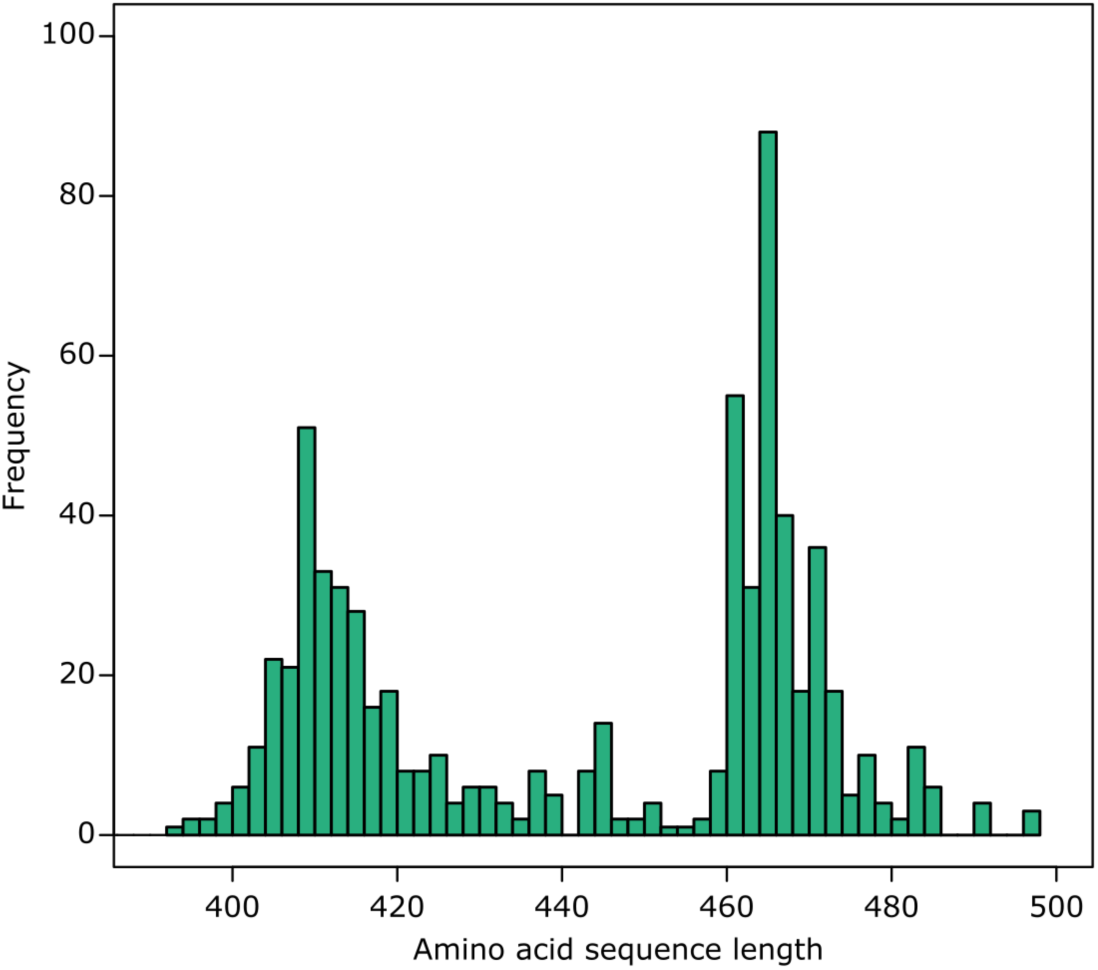
XusB BLAST hit amino acid sequence length distribution. The BtXusB amino acid sequence was submitted to the EFI Enzyme Similarity Tool server (*6*). BLAST results were filtered to remove hits with an E-value higher than 10^-5^ and sequences shorter than 380 and longer than 500 amino acids, resulting in 681 hits. The amino acid sequence lengths of these hits are plotted in the histogram, clearly showing a bimodal distribution with peaks around 410 and 465 amino acids. The shorter group includes BfXusB, while BtXusB, BvXusB and *P. distasonis* XusB belong to the longer group.

**Fig. S10.**
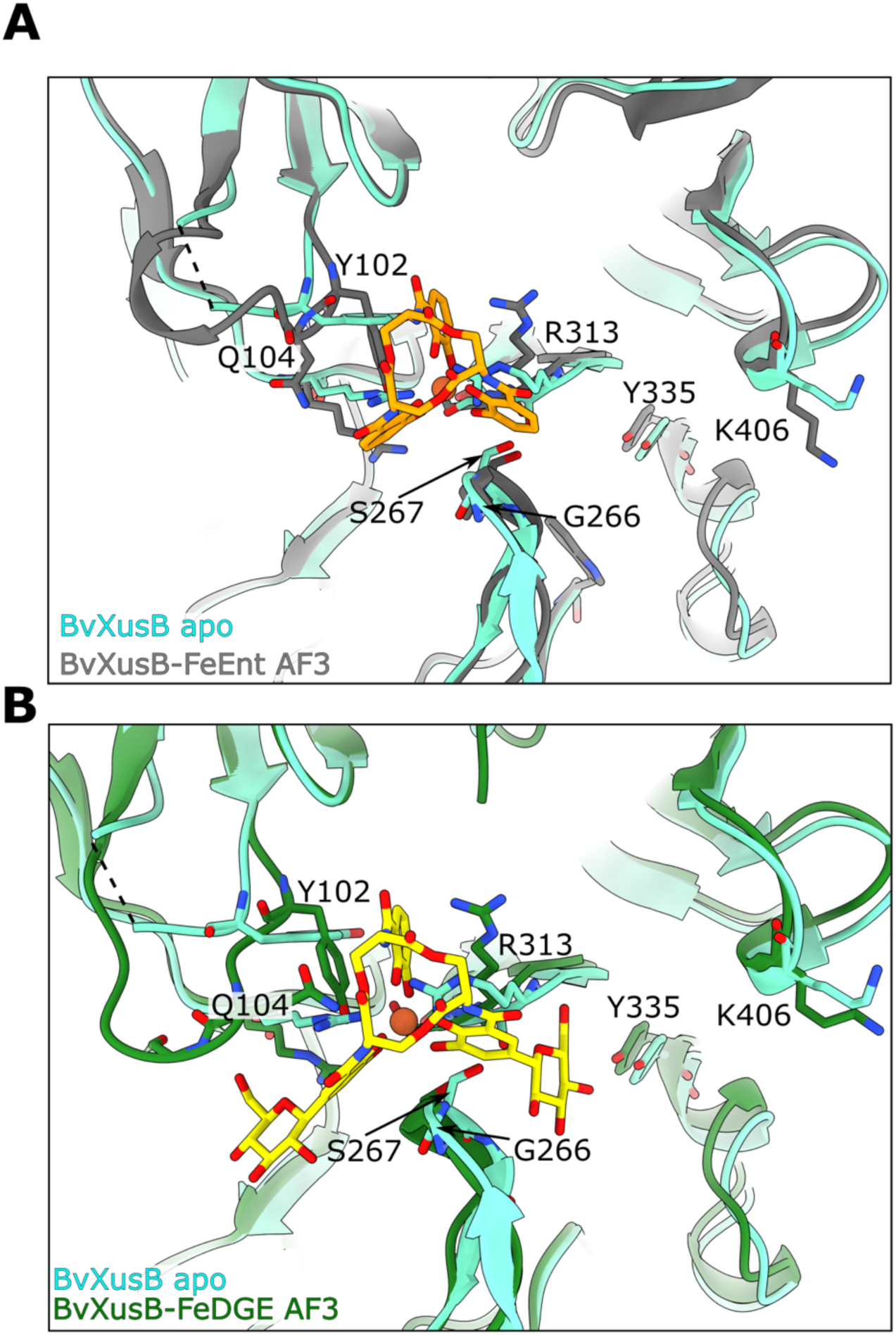
Comparison of BvXusB apo structure and ligand-bound structure predictions. BvXusB apo crystal structure (cyan) superposed with AF3-predicted BvXusB-FeEnt (**A**) and BvXusB-FeDGE (**B**) models.

**Fig. S11.**
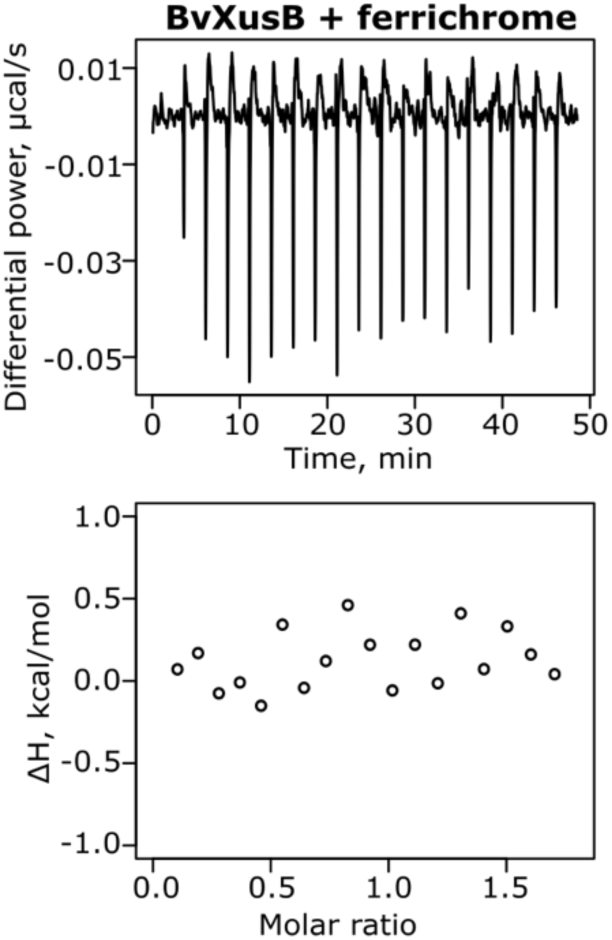
BvXusB does not bind ferrichrome. Representative ITC experiment where 250 μM ferrichrome was titrated into 25 μM BvXusB (n=2). No substantial injection heats were observed, consistent with lack of binding.

**Table S1.**
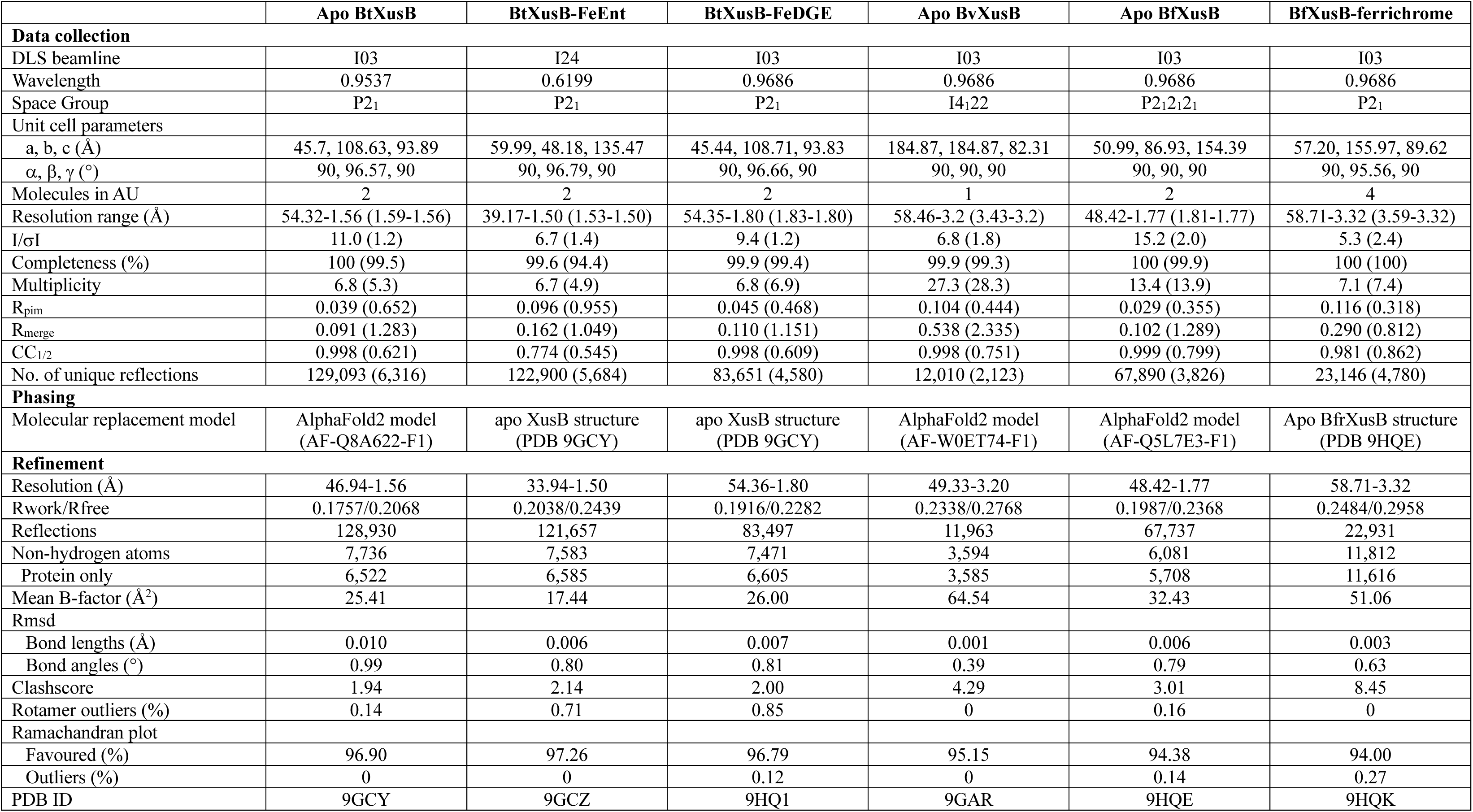
X-ray data collection and refinement parameters. Values for highest resolution shell in parentheses.

**Table S2.** Isothermal titration calorimetry data fitting results.

| Titration | Replicate | [Protein] (M) | [Ligand] (M) | N | Kd (M) | $\Delta H$ (kcal/mol) | $\Delta G$ (kcal/mol) | $-T\Delta S$ (kcal/mol) |
| --- | --- | --- | --- | --- | --- | --- | --- | --- |
| BtXusB + FeEnt | 1 | 2.50E-04 | 2.50E-05 | 1.3 | 9.82E-08 | -24.5 | -9.57 | 14.9 |
| BtXusB + FeEnt | 2 | 2.50E-04 | 2.50E-05 | 1.46 | 8.42E-08 | -22.8 | -9.67 | 13.1 |
| BtXusB + FeEnt | 3 | 2.50E-04 | 2.50E-05 | 1.47 | 4.94E-08 | -21 | -9.97 | 11 |
| BtXusB + FeEnt | 4 | 2.50E-04 | 2.50E-05 | 1.22 | 5.52E-08 | -22.6 | -9.9 | 12.7 |
| BtXusB + FeDGE | 1 | 1.54E-04 | 1.73E-05 | 1 (fixed) | 1.69E-07 | -20.1 | -9.24 | 10.8 |
| BtXusB + FeDGE | 2 | 1.57E-04 | 1.73E-05 | 1 (fixed) | 1.99E-07 | -19.5 | -9.14 | 10.4 |
| BvXusB + FeEnt | 1 | 2.50E-04 | 2.50E-05 | 1.35 | 1.01E-07 | -15.5 | -9.55 | 5.98 |
| BvXusB + FeEnt | 2 | 2.50E-04 | 2.50E-05 | 1.21 | 7.87E-08 | -14.8 | -9.69 | 5.15 |
| BfXusB + ferrichrome | 1 | 2.50E-04 | 2.50E-05 | 1.18 | 2.68E-07 | -11.5 | -8.97 | 2.52 |
| BfXusB + ferrichrome | 2 | 3.48E-04 | 2.54E-05 | 1.27 | 2.99E-07 | -10.1 | -8.9 | 1.24 |
| BfXusB + ferrichrome | 3 | 2.46E-04 | 2.54E-05 | 1.09 | 1.76E-07 | -10.3 | -9.22 | 1.11 |
| BfXusB + ferrichrome | 4 | 2.26E-04 | 2.27E-05 | 1.06 | 2.98E-07 | -12.7 | -8.9 | 3.78 |

**Table S3.** Cryo-EM data collection and structure refinement parameters.

|  | **XusAB** |
| --- | --- |
| **Data collection** |  |
| Electron microscope | FEI Titan Krios |
| Voltage (kV) | 300 |
| Spherical aberration (μm) | 2.7 |
| Camera | Falcon 4i (counting) |
| Energy filter | Selectris (10 eV slit) |
| Magnification | 165,000 |
| Pixel size (Å) | 0.74 |
| Total dose (e<sup>-</sup>/Å<sup>2</sup>) | 40.83 |
| Defocus range (μm) | -2.0 to -0.8 |
| Number of movies collected | 6,645 |
| **Image Processing** |  |
| Symmetry | C1 |
| Initial number of particles | 1,110,843 |
| Final number of particles | 76,969 |
| Global resolution (FSC = 0.143) | 2.7 Å |
| Map sharpening B-factor (Å<sup>2</sup>) | -57.2 |
| **Refinement** |  |
| Model composition |  |
| Non-hydrogen atoms | 8,742 |
| Protein residues | 1,093 |
| R.m.s. deviations |  |
| Bonds lengths (Å) | 0.002 |
| Bond angles (°) | 0.543 |
| Validation |  |
| MolProbity score | 1.12 |
| Clash score | 2.84 |
| Rotamer outliers (%) | 0 |
| Ramachandran plot |  |
| Favoured (%) | 97.79 |
| Outliers (%) | 0 |
| PDB | 9GBC |
| EMDB | EMD-51210 |

**Table S4.**
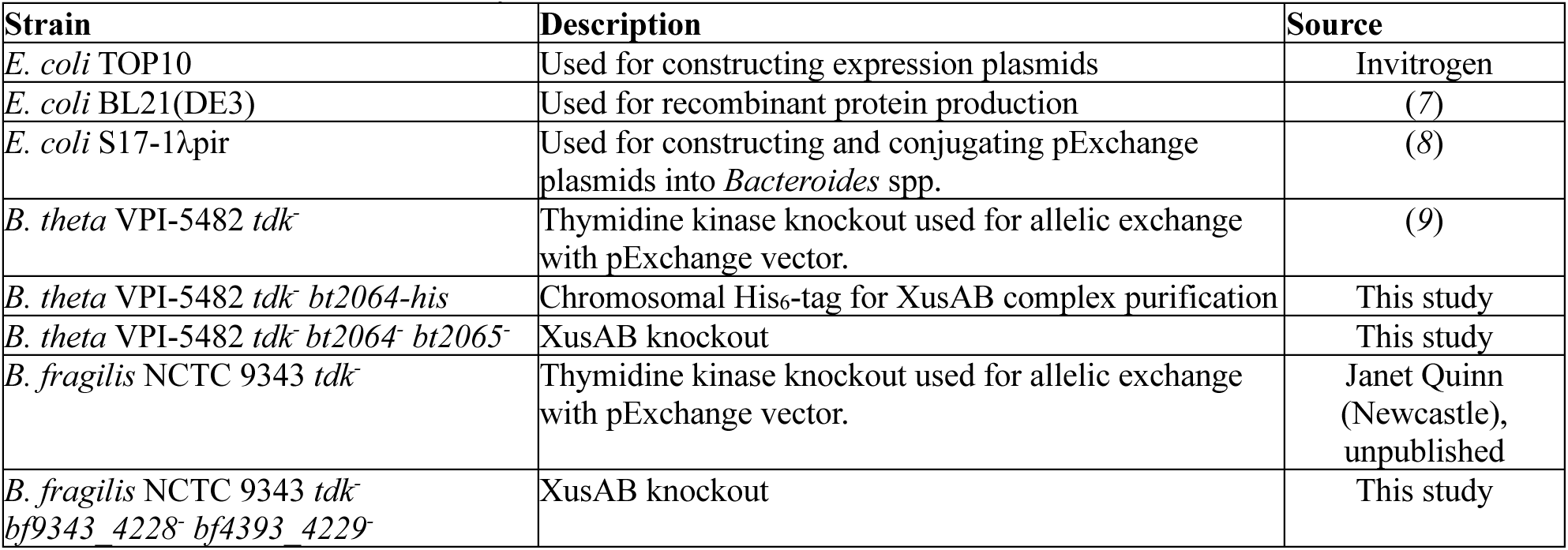
Strains used in this study.

**Table S5.**
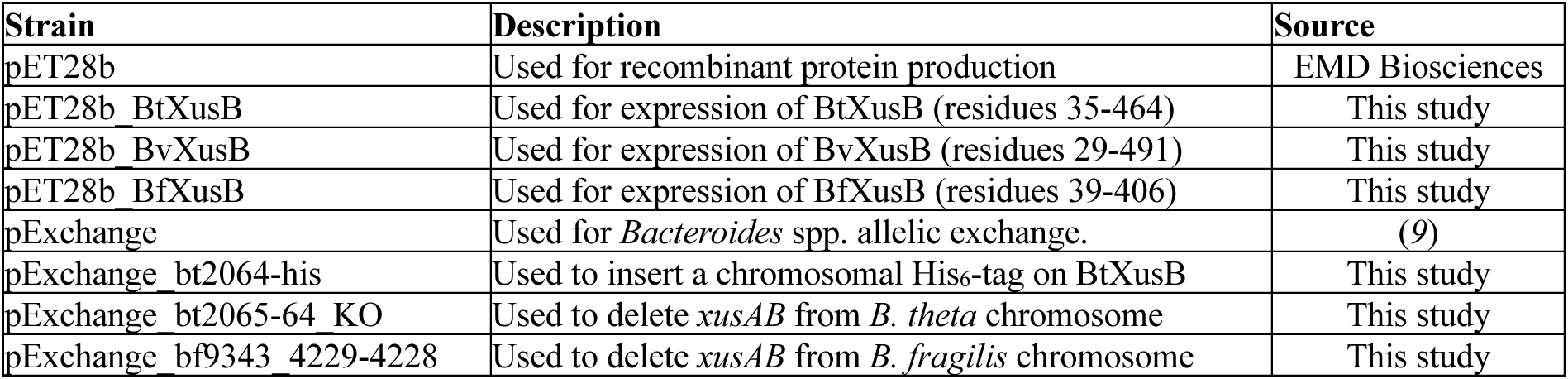
Plasmids used in this study.

**Movie S1.** Morph of the apo BtXusB crystal structure to the FeEnt-bound BtXusB crystal structure. The protein model is depicted as a cartoon and as a surface. The siderophore-binding loops are in blue. The position of FeEnt, shown as an orange stick model, is fixed for reference.

**Movie S2.** Fit of FeEnt and the BtXusB residues and interacting water molecules to the 2mF_o_-DF_c_ electron density map contoured at 1.5σ. FeEnt is in orange, BtXusB residues are in grey, and water molecules are shown as red spheres.

